## Supplemental (Apendix) Figures S1-S33 for "Dormancy-to-death transition in yeast spores occurs due to gradual loss of gene-expressing ability"

---

### Table of contents:

**- Appendix Figures S1-S33** ........................................................................ **Pgs. 2 - 50**

**- Appendix Table S1** ........................................................................ **Pgs. 51 - 53**

### Appendix Figures

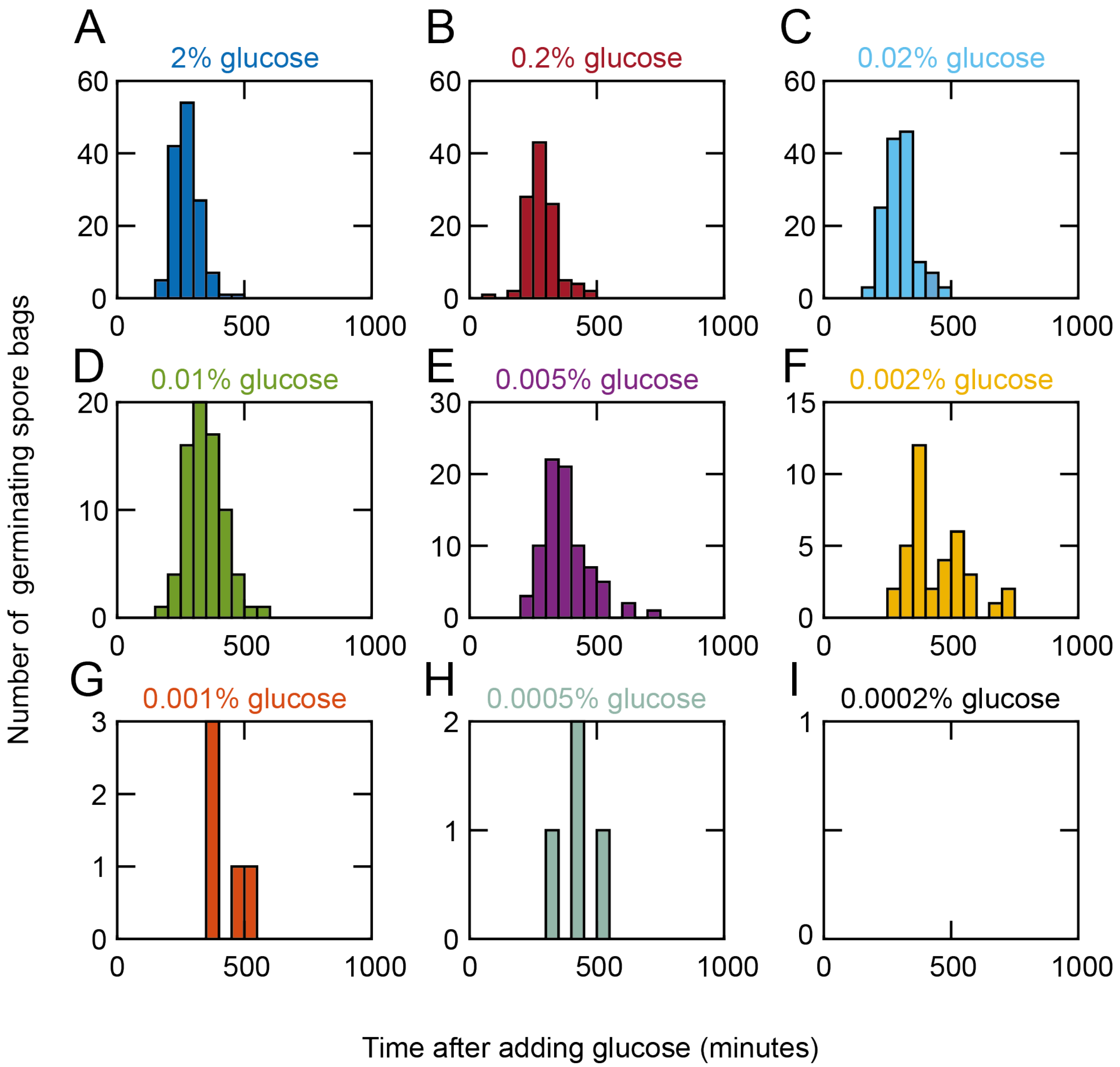


**Appendix Figure S1 - Representative histograms showing how many wild-type spore bags typically germinated for each glucose concentration (complements Figs. 1B-C)**. We incubated the spores in minimal media (which contains all essential amino acids and nitrogenous bases) that we supplemented with a desired concentration of glucose (denoted above each histogram). We then used a wide-field microscope to observe individual spore bags over time and noted how many spore bags germinated as a function of time after we added glucose. For each glucose concentration, the number of spore bags that we analyzed for the histograms shown here are: **(A)** 2%-glucose, *n* = 137 spore bags; **(B)** 0.2%-glucose, *n* = 117 spore bags; **(C)** 0.02%-glucose, *n* = 142 spore bags; **(D)** 0.01%-glucose, *n* = 78 spore bags; **(E)** 0.005%-glucose, *n* =

103 spore bags; **(F)** 0.002%-glucose, *n* = 190 spore bags; **(G)** 0.001%-glucose, *n* = 106 spore bags; **(H)** 0.0005%-glucose, *n* = 95 spore bags; **(I)** 0.0002%-glucose, *n* = 50 spore bags (note that none of these spore bags germinated at this glucose concentration).

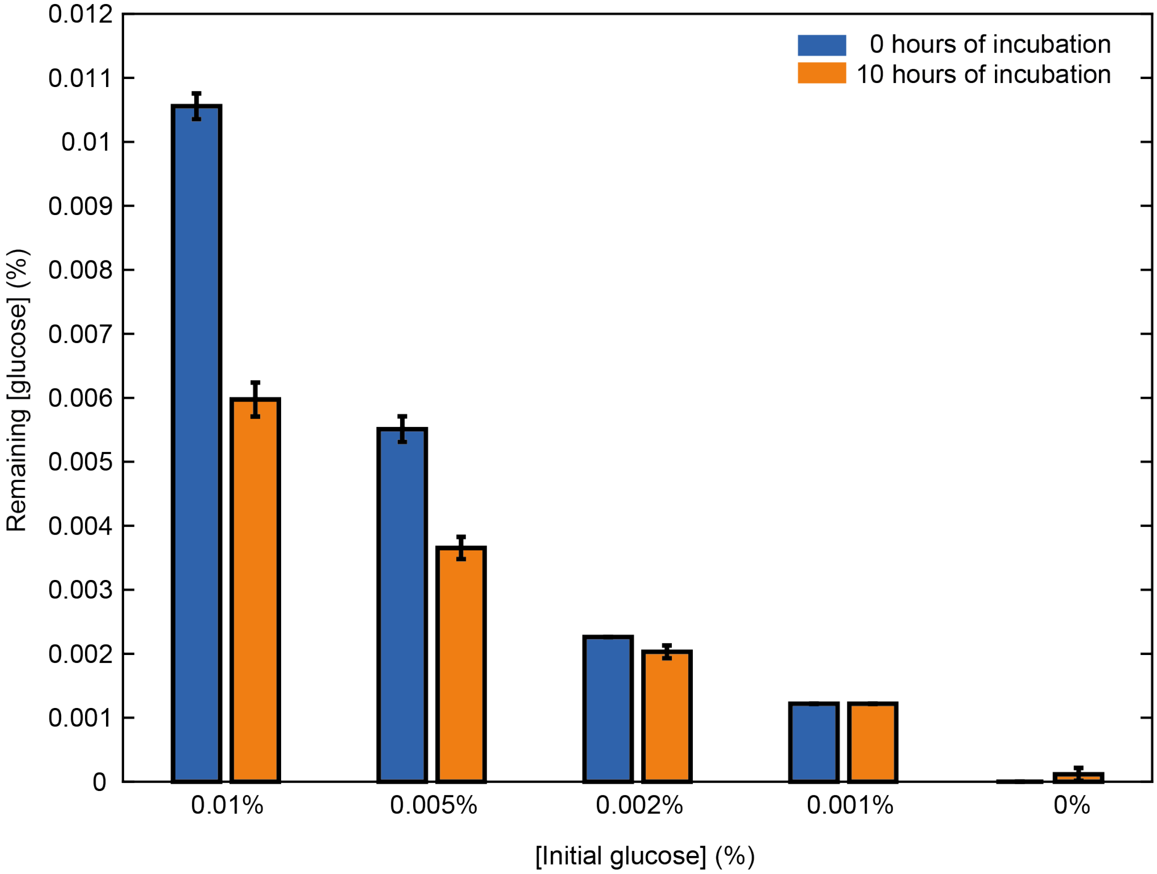


**Appendix Figure S2 - Running out of glucose is not the reason for only a fraction of spore bags germinating for non-saturating glucose-concentrations (complements Figs. 1C-D).** Before incubating the wild-type spores in a minimal medium with glucose (experiment shown in Fig. 1C), we measured the initial glucose-concentration in the medium (blue bars - "0 hours of incubation") with a hexokinase-based assay (see Methods). In all the glucose concentrations that we studied (from 0.0002% to 2%), all germinations have either stopped or were about to stop after ~10 hours (~600 minutes) of incubation. This is evidenced by the plateauing of all the curves - representing the % of spore bags germinated as a function of time for various initial glucose-concentrations - beginning at around 600 to 700 minutes in Fig. 1C. We sought to determine whether the germinations were stopping due to vegetative cells, which result from germinated spores, having consumed appreciable amounts of glucose during their continuous cell divisions (vegetative yeasts divided once every 2-3 hours at these glucose concentrations). We measured the remaining glucose-concentration after ~10 hours of incubation for various initial glucose-concentrations (orange bars - "10 hours of incubation"). For the relatively high initial glucose-concentrations (e.g., 0.01% and 0.005% shown here), germinated spores and the resulting vegetative cells depleted nearly half of the initial glucose-concentration after 10 hours. But for the relatively low initial glucose-concentrations (e.g., 0.002% and 0.001% shown here), the germinated spores and the resulting vegetative cells have not consumed appreciable amounts of glucose - the glucose-concentration remained nearly unchanged after 10 hours. These results show that the germinations do not stop at any of the glucose-concentrations that we studied (Fig. 1C) because the spores ran out of glucose. This is also true for the relatively high glucose-concentrations (e.g., 0.01% and 0.005%) since these conditions still had high amounts of glucose remaining after 10 hours - these remaining glucose-concentrations were still higher than some the low initial glucose-concentrations (e.g., 0.002% and 0.001%) which were enough to germinate spores. For all bars shown: $n$ = 3, error bars are s.e.m.

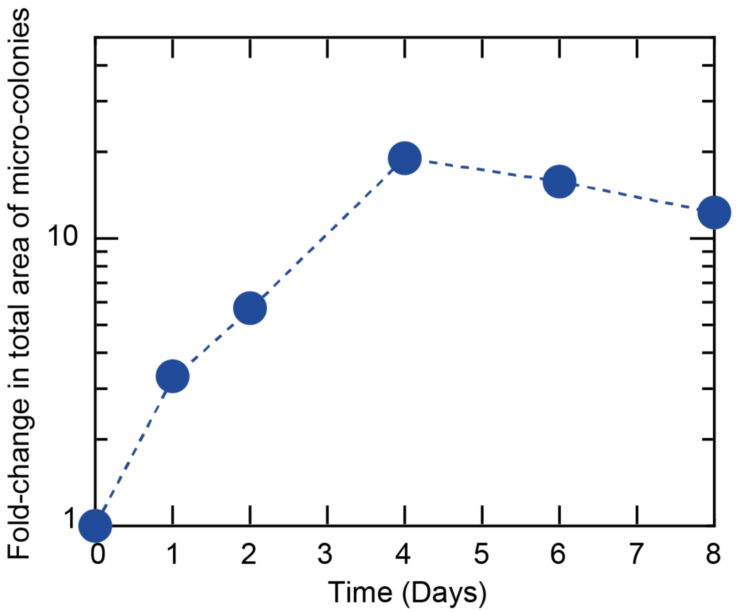


**Appendix Figure S3 - A vegetative yeast cell can divide multiple times even with the lowest glucose-concentration (0.0002%) that we used, which could not germinate any spores (complements Figs. 1C-D).** We sought to test if the lowest concentration of glucose that we used in Figs. 1C-D, which was 0.0002% and could not germinate any spores, was enough to allow divisions of vegetative, diploid wild-type cells that formed the spore bags. We incubated the vegetative, diploid wild-type cells in a minimal medium with 0.0002% glucose and used a wide-field microscope to observe them over 8 days. We found that these cells could divide multiple times. Specifically, we took pictures of 10 micro-colonies on different days and measured the area of each micro-colony over time. We then combined all their areas into a single number, on each day, and thus determined the fold-change in the total (combined) area of the colonies over time (i.e., the combined area of all micro-colonies on each day divided by the combined area of all the micro-colonies on day 0). On the fourth day, the colonies stopped growing, at which point the total area had increased by ~19-fold compared to the initial total area (corresponds to ~4.2 divisions). As a control, we incubated the vegetative cells in a minimal medium without any glucose, which led to a lesser, transient growth that stopped after two days (data not shown). These results establish that 0.0002% of glucose is enough to sustain multiple divisions of a single, isolated, vegetative wild-type cell despite it not germinating any spores.

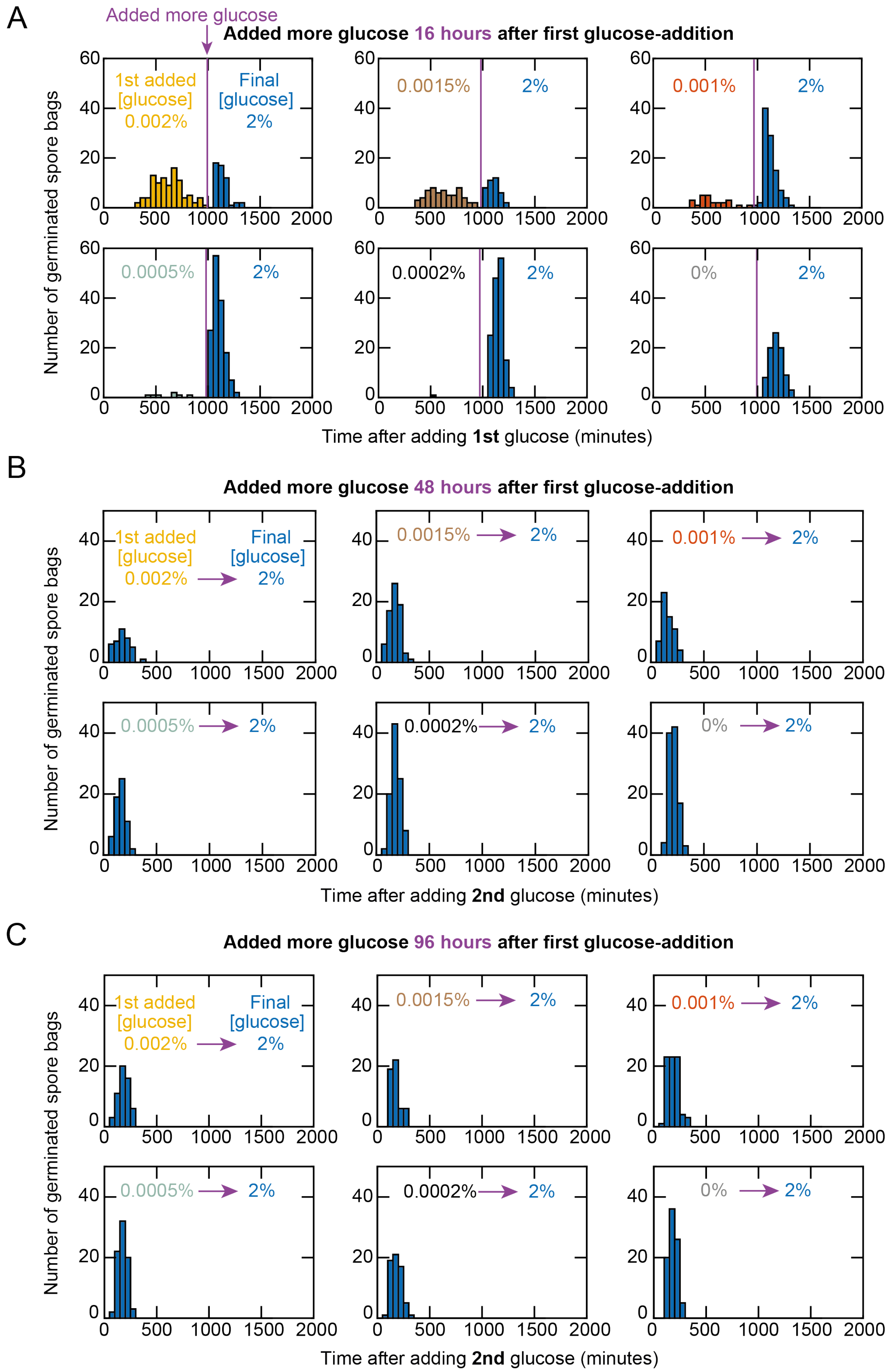


**Appendix Figure S4 - Representative histograms showing how many wild-type spore-bags typically germinate when we add glucose in two steps as shown in Fig. 2A (complements**

**Fig. 2B).** In the experiment described in Fig. 2A, we first incubated the wild-type spores in a minimal medium with a relatively low concentration of glucose. We used a wide-field microscope to observe the spores and count the number of spore bags that germinated as a function of time. A set number of hours after incubating the spores in the low glucose-concentration (16 hours in **(A)**, 48 hours in **(B)**, and 96 hours in **(C)**), we added more glucose to the medium and then counted how many more spore bags germinated as a result - these spore bags did not germinate when we gave them the first, low concentration of glucose. Shown here are typical histograms from these experiments. **(A)** Adding more glucose after 16 hours (~1000 minutes) of incubation in the lower glucose-concentration (indicated within each of the histograms) so that, after adding the second batch of glucose, the final glucose-concentration was 2% (saturating concentration). In each histogram, the colored bars that appear to the left of the purple vertical line show germinations that occur in the first glucose-concentration while the blue bars that are to the right of the purple vertical line show germinations that occur after adding more glucose. Time here is the time after adding the first batch of glucose. The number of spore bags that we analyzed for each histogram are (including those that did not germinate and thus not shown as bars in the histograms): *n* = 208 (top left); *n* = 126 (top center); *n* = 126 (top right); *n* = 220 (bottom left); *n* = 150 (bottom center); *n* = 100 (bottom right). **(B)** Showing histograms only for the germinations that occur after adding the second batch of glucose (48 hours (2 days) after adding the first batch of glucose) so that the final glucose-concentration was 2%. Time here is the time after adding the second batch of glucose. We do not show the germinations that occur before adding the second batch of glucose because they resemble the data show in (A). The first glucose-concentrations are indicated in each histogram. The number of spore bags that we analyzed for each histogram are (including those that did not germinate and thus not shown as bars in the histograms): *n* = 41 (top left); *n* = 49 (top center); *n* = 60 (top right); *n* = 63 (bottom left); *n* = 98 (bottom center); *n* = 106 (bottom right). **(C)** Showing histograms only for the germinations that occur after adding the second batch of glucose (96 hours (4 days) after adding the first batch of glucose) so that the final glucose-concentration was 2%. Time here is the time after adding the second batch of glucose. We do not show the germinations that occur before adding the second batch of glucose because they resemble the data show in (A). The first glucose-concentrations are indicated in each histogram. The number of spore bags that we analyzed for each histogram are (including those that did not germinate and thus not shown as bars in the histograms): *n* = 58 (top left), *n* = 53 (top center); *n* = 77 (top right); *n* = 79 (bottom left); *n* = 65 (bottom center); *n* = 88 (bottom right).

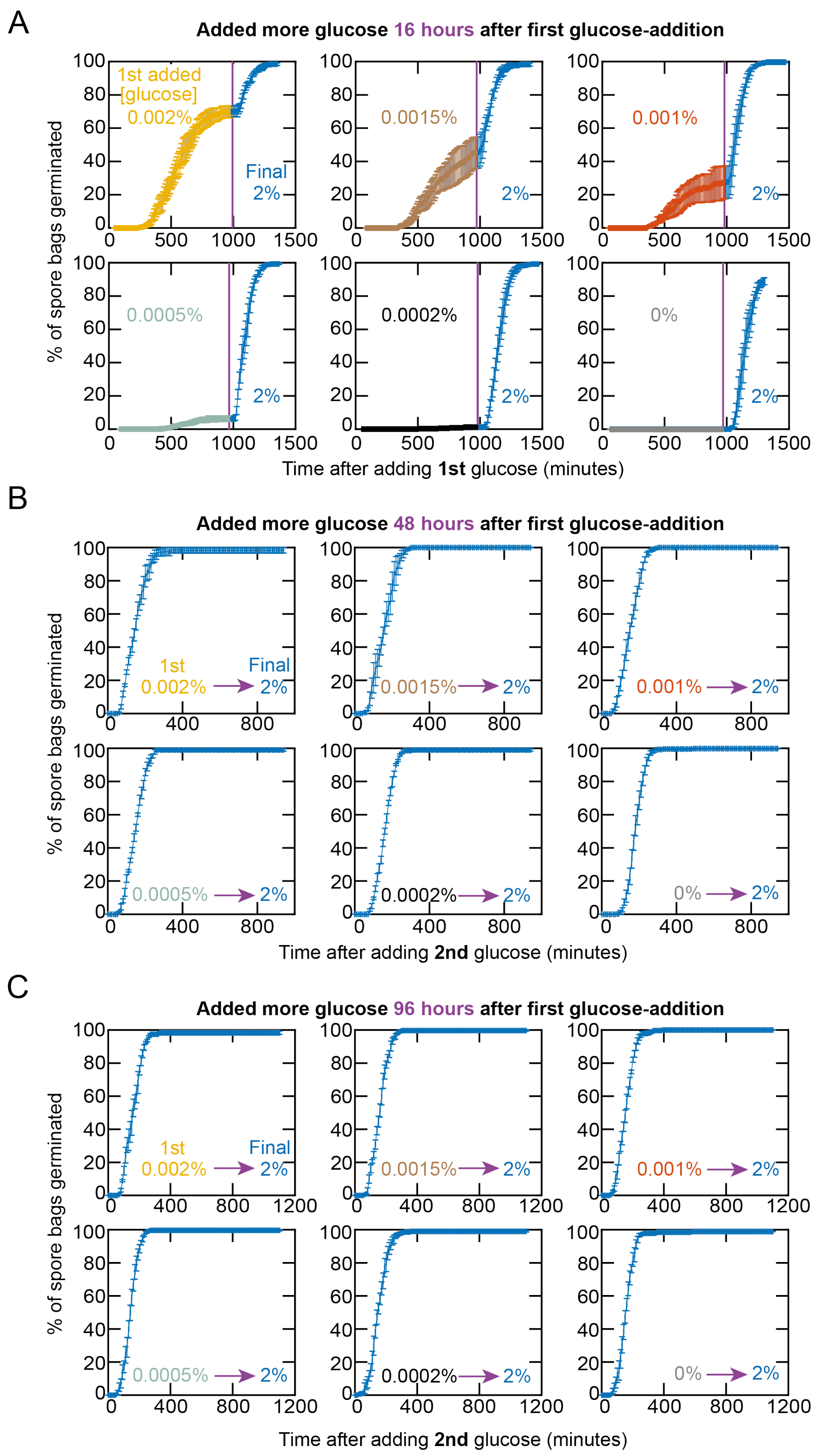


**Appendix Figure S5 - Percentage of wild-type spore-bags that germinated as a function of time in experiments in which we add glucose in two steps as shown in Fig. 2A (summarizes**

**data shown in Appendix Fig. S4 and complements Fig. 2C). (A)** Added more glucose after 16 hours (~1000 minutes) of incubation in the lower glucose-concentration (indicated within each of the histograms) so that, after adding the second batch of glucose, the final glucose-concentration was 2% (saturating concentration). Summarizes the histograms shown in Appendix Fig. S4A by plotting the percentage of the wild-type spore-bags that have germinated after some time (i.e., cumulative percentage of germinations as a function of time). Time here is the time after adding the first batch of glucose. Purple line indicates when we added the second batch of glucose that raised the glucose-concentration to a saturating value (2%). The first glucose-concentration (to the left of the purple vertical line) is indicated in each histogram. **(B)** Summarizes the histograms shown in Appendix Fig. 4B. Data shown only for the germinations that occur after adding the second batch of glucose (48 hours (2 days) after adding the first batch of glucose) so that the final glucose-concentration was 2%. We do not show the germinations that occur before adding the second batch of glucose because they resemble the data show in (A). Time here is the time after adding the second batch of glucose. **(C)** Summarizes the histograms shown in Appendix Fig. 4C. Data shown only for the germinations that occur after adding the second batch of glucose (96 hours (4 days) after adding the first batch of glucose) so that the final glucose-concentration was 2%. We do not show the germinations that occur before adding the second batch of glucose because they resemble the data show in (A). Time here is the time after adding the second batch of glucose. In all plots (A-C): $n$ = 3 and error bars are s.e.m.

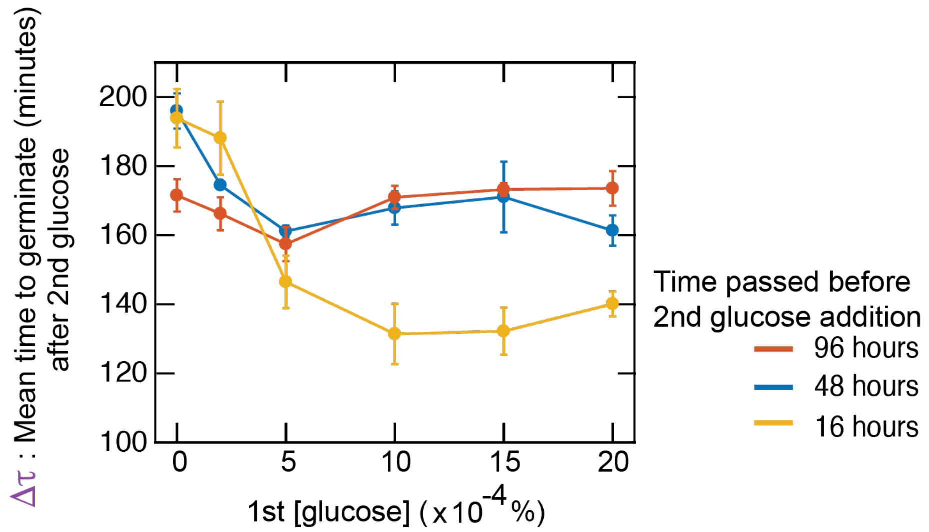


**Appendix Figure S6 - Average time taken by wild-type spore-bags to germinate, as a function of the first, low glucose-concentration that primes the spores, in experiments in which we add glucose in two steps (shown in Fig. 2A) (complements Fig. 2F).** We gave glucose in two steps to wild-type spore-bags as indicated in Fig. 2a. We first added a relatively low glucose-concentration (indicated along the horizontal axis in the plot). After 16 hours (yellow) or 48 hours (blue) or 96 hours (red), we added more glucose to raise the total glucose-concentration to 2% (saturating level). The mean time ($\Delta\tau$) plotted along the vertical axis represents the time after adding the second batch of glucose. These are the same data as the ones plotted in Fig. 2F but now shown in different units - the average time ($\Delta\tau$) is now in minutes whereas Fig. 2F shows, for each color, the $\Delta\tau$ after dividing it by the average time taken by the spore bags that did not see any glucose (0%-glucose condition) before receiving the second batch of glucose. $n$ = 3; error bars are s.e.m.

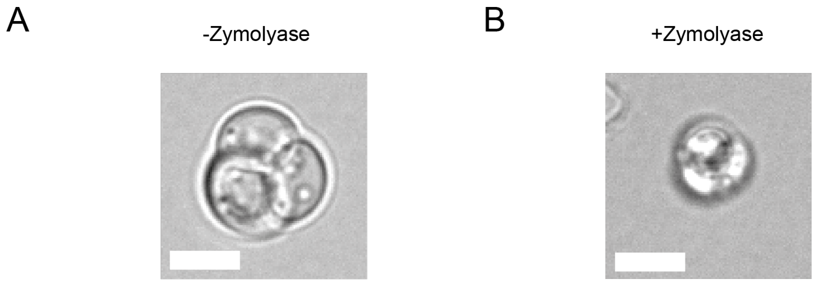


**Appendix Figure S7 - Zymolyase leaves un-germinated spore-bags intact while lysing vegetative yeasts that result from germinated spores (complements Fig. 2G).** In a mixture of spore-bags and vegetative yeasts, one typically isolates the spore bags by using zymolyase, which lyses vegetative yeasts but not spore bags due to the spore bags' thick, protective outer walls. For this reason, typical (but not all) sporulation procedures (i.e., procedures for forming spores from diploid yeasts) involve adding zymolyase at the end to isolate spore bags and kill off any diploid yeasts that failed to form spore bags. We did not use zymolyase at the end of our sporulation procedure because we typically had high yields of spore bags and, more importantly, zymolyase hurts the spore bags by causing them to lose their protective walls (seen in (B)). We did not want to hurt the spore bags in our experiments. Since our experiments involved using a microscope to track individual spore bags, we could always distinguish vegetative cells from spores. Thus, we did not need to add zymolyase at the end of our sporulation procedure (zymolyase is necessary for population-level experiments in which one does not track individual spore bags). But we used zymolyase to isolate primed, un-germinated spores (Fig. 2E) from the vegetative cells that resulted from the spore bags that did germinate. To be sure, we checked by microscopy that zymolyase indeed lysed vegetative cells and left behind only un-germinated spore bags. **(A)** A representative microscope-image that shows un-germinated spore bags in the absence of zymolyase (scale bar, 5 μm). **(B)** A representative microscope-image that shows intact, un-germinated spore bags after the zymolyase treatment (not the same field of view as (A)). As seen here, spore bags appear smaller after encountering zymolyase than they did before they encountered zymolyase because zymolyase partially degrades their protective walls (note the lack of white outline in (B) that exists around the spore bag in (A). But the spores are still intact and kept together as one unit inside a bag, as seen here (scale bar, 5 μm). After adding zymolyase, we immediately proceeded to the next step, in which we lysed the spores to extract their RNAs for RNA-seq (Fig. 2G and Appendix Fig. S8).

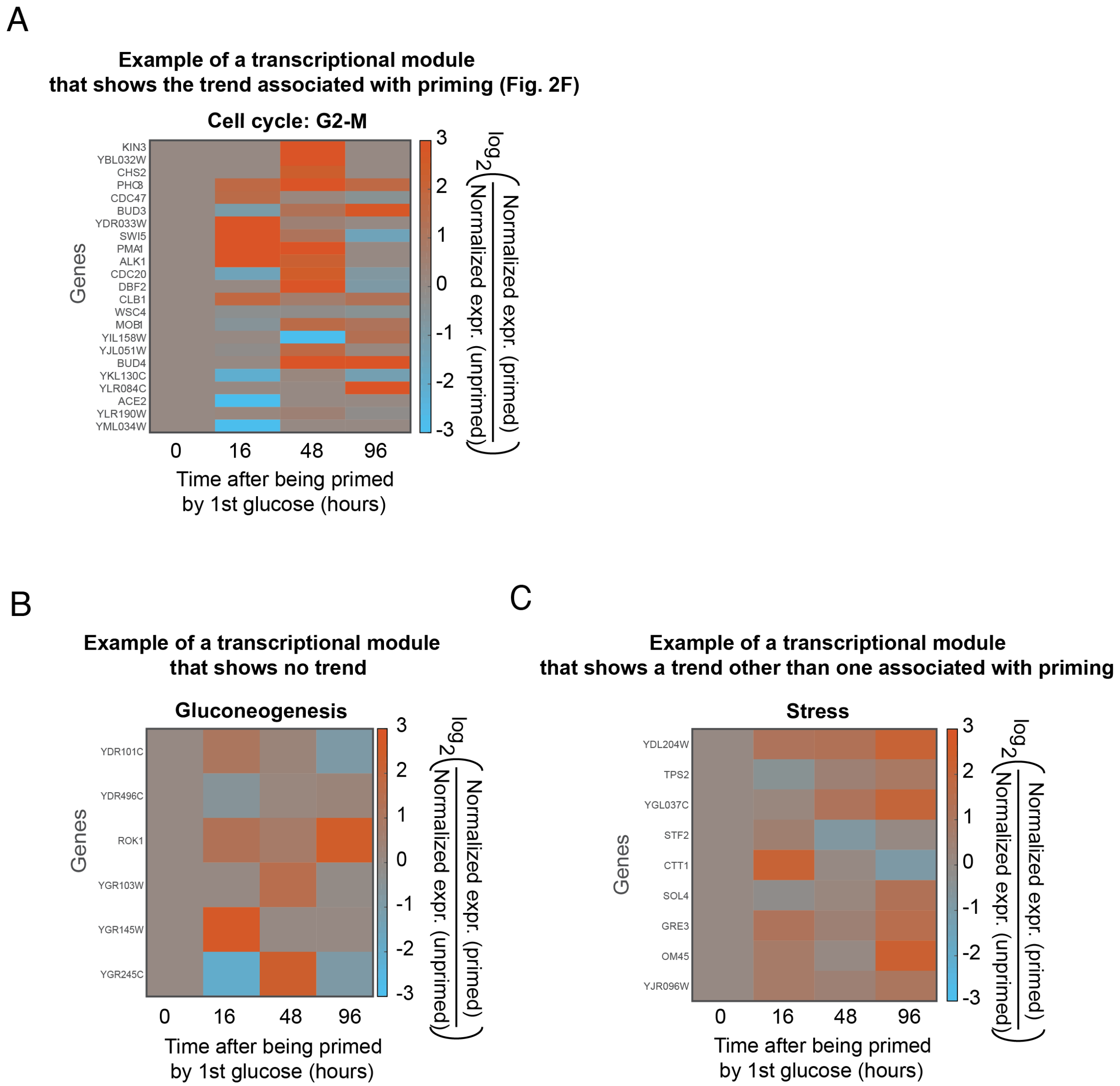


**Appendix Figure S8 - Genome-wide view of primed dormancy (representative transcriptional modules shown; complements Fig. 2G).** As described in the main text and in Materials and Methods, we performed a transcriptome (RNA-Seq) analysis of spores that did not germinate after encountering a low glucose-concentration. This plot showcases details of RNA-Seq analyses that led to the summarized results shown in Fig. 2G. Our analysis relies on an insightful previous work by Joseph-Strauss et al. (2007) (*10*) that identified a list of transcriptional modules by studying yeast spores that germinated after receiving a saturating concentration (2%) of glucose (list of genes that they found for each transcriptional module, which we used here, is in Appendix Table S1). In the three heat maps shown here, the colors represent normalized gene-expression levels for individual genes within each transcriptional module whereas Fig. 2G shows a single, normalized gene-expression for an entire transcriptional module at each time point that we obtained by averaging the expression levels of all genes in that module. "Normalization" for (A-C) means that we divided the expression level of a given gene for spores that received a low concentration of glucose - which did not germinate them - by the expression level of the same gene for spores that were kept in minimal media without glucose for the same amount of time as the spores that received the glucose (for Fig. 2G, we normalized in the same way except that we used the average expression level of a module instead of individual genes). Here we show representative transcriptional modules that reveal how studying individual genes can give a different perspective from the one provided by averaging over all genes in a module (Fig. 2G). **(A)** Normalized gene-expression profiles of the “Cell-cycle: G2-M” module (list of genes in Appendix Table S1). When we average the expression levels of all genes in this module for each time point (Fig. 2G), we observe a clear temporal trend that qualitatively mirrors the temporal trend in the average time taken by the primed spores to germinate (Fig. 2F) - namely, the normalized gene-expression level for the module (Fig. 2G) is elevated after 16 hours and 48 hours but decays away after 96 hours. Interestingly, we observe the same trend for a number of individual genes in this module, with slight differences in timing (i.e., heat map here shows some genes having their expression level peaking at 16 hours while others do so at 48 hours). **(B)** Normalized gene-expression profiles of the “gluconeogenesis” module (list of genes in Appendix Table S1). When we average the expression levels of all genes in this module for each time point (Fig. 2G), we observe no clear trend. However, when we study the expression levels of individual genes in this module, as shown here, we observe diverging trends (i.e., some genes have a red pixel while others have a green pixel at the same time point), explaining the absence of any observable trends when we average over all genes to get a single expression-level for this module (Fig. 2G). **(C)** Normalized gene-expression profiles of the “stress” module (list of genes in Appendix Table S1). When we average the expression levels of all genes in this module for each time point (Fig. 2G), we observe a clear trend (i.e., elevated expression-level over time) that is qualitatively different from the temporal trend in the average time taken by the primed spores to germinate (Fig. 2F). When we study the expression levels of individual genes in this module, as shown here, we see a homogeneous expression profile (i.e., all genes have red pixels at and after 16 hours - no temporal undulations in the expression levels over time).

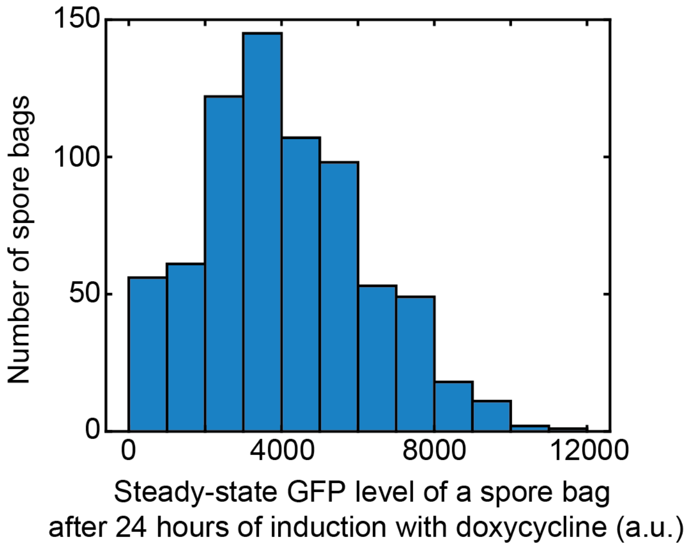


**Appendix Figure S9 - GFP expression by dormant spores without any nutrients (in PBS) (complements Fig. 3C).** We incubated a population of GFP-inducible spores (Fig. 3A) in a saline solution (PBS) with 100 µg/ml of doxycycline for 24 hours. By the end of the 24-hour incubation, spore bags' GFP levels reach steady-state values (Fig. 3C). At the end of the 24-hour incubation, we measured the steady-state GFP level of each spore bag in the population with a wide-field, epifluorescence microscope and plotted the distribution of their GFP levels here ($n$ = 723 spore bags counted). To measure the GFP level of a single spore bag, we computed the average intensity of all the pixels that belonged to a single spore bag in a microscope image in the GFP-channel. This average is the steady-state GFP level of a spore bag which we plotted here for multiple spore bags in a population. Here, we also subtracted the background fluorescence value so that a GFP level of zero represents a spore bag with no GFP (i.e., it has the same fluorescence as the background).

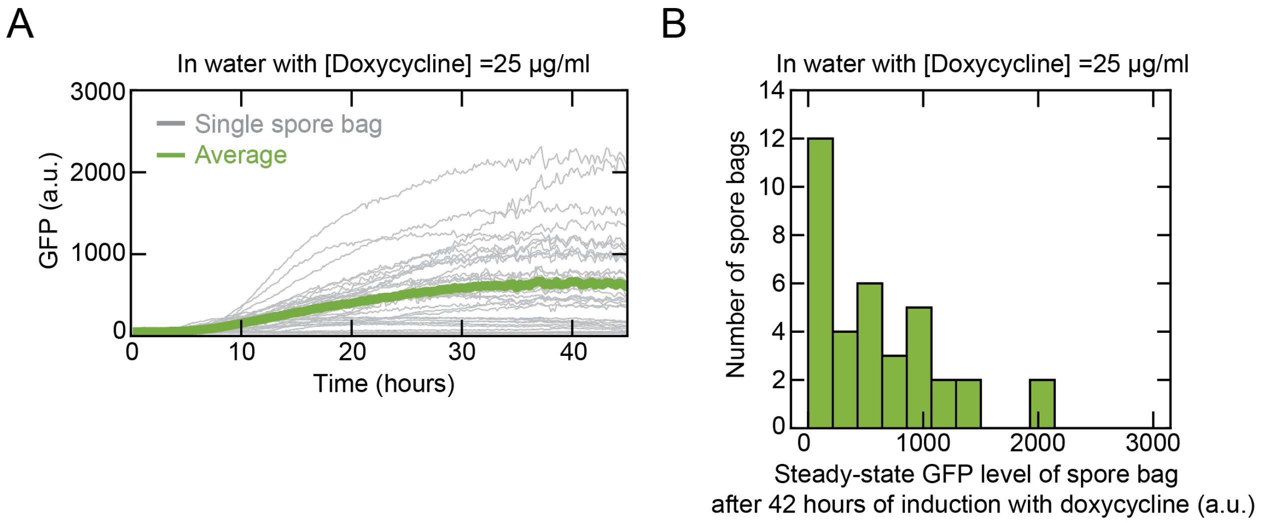


**Appendix Figure S10 - Induction of GFP expression is possible for spores that are in water without any nutrients (no amino acids and no glucose) (complements Fig. 3B). (A)** GFP level of individual spore bags (grey curves) over time during 42 hours of incubation in Milli-Q water (dd$H_2O$) with 25 µg/ml of doxycycline (*n* = 36 spore bags). Green curve is the GFP-level averaged over all the spore bags. We see here that the spore bags require about 30 to 40 hours to produce steady-state levels of GFP in water whereas they require ~20 hours to do so in PBS with the same doxycycline concentration (see Appendix Fig. S12C). **(B)** Steady-state GFP-levels of individual spore bags after 42 hours of incubation in Milli-Q water with 25 µg/ml of doxycycline. These results show that spores, in plain water without any nutrients (no amino acids and no glucose), can still highly express GFP - as high as vegetative yeasts (compare (B) with Appendix Fig. S11C). Interestingly, while spore bags in PBS reach half of their saturating GFP levels after 10 hours of induction (see Fig. 3C or Appendix Fig. S12), we see here that the spore bags in water take ~30 to 40 hours to reach steady-state GFP levels. In particular, as seen in (A), spore bags in water have nearly undetectable levels of GFP even ~10 hours after encountering doxycycline. On the other hand, for the same doxycycline concentration, the average steady-state GFP-levels are similar between spore bags in water and spore bags in PBS.

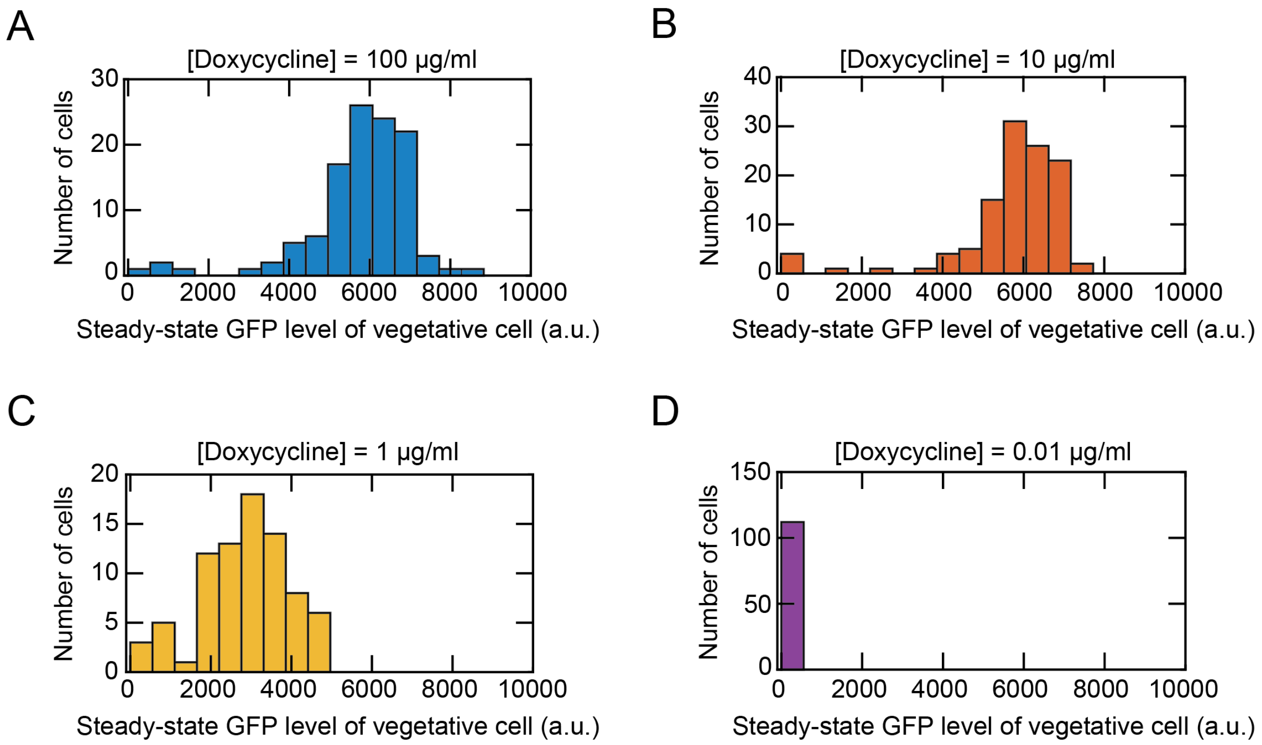


**Appendix Figure S11 - Steady-state GFP levels of vegetative (replicating) cells that have the same synthetic gene-circuit as the GFP-inducible spores (Fig. 3A) (for comparison with the steady-state GFP levels of spore bags) (complements Fig. 3C).** We sought to compare how the GFP levels of spores compare with the GFP levels of replicating cells that have the same gene circuit. Shown here are the steady-state GFP-levels of individual, replicating, diploid cells that have the same GFP-inducing gene-circuit as the GFP-inducible spores (Fig. 3A). In fact, we sporulated these cells to form the GFP-inducible spores. We measured these GFP levels after 8 hours of incubation in minimal media with a 2%-glucose and **(A)** 100 µg/ml of doxycycline (*n* = 112 cells), or **(B)** 10 µg/ml of doxycycline (*n* = 113 cells), or **(C)** 1 µg/ml of doxycycline (*n* = 80 cells), or **(D)** 0.01 µg/ml of doxycycline (*n* = 108 cells). Data obtained by using wide-field epifluorescence microscopy as in Appendix Fig. S9. By comparing these GFP levels of diploid, replicating cells with the GFP levels of spores (Appendix Fig. S9), we see that even without nutrients, spores can produce GFP at levels that are similar to those of replicating cells. The two main differences between the spores and vegetative cells is that (1) the spores without nutrients requires more time (~24 hours) to reach steady-state GFP levels where as the vegetative cells require only ~ 8 hours to reach steady-state GFP levels and that (2) spores need more doxycycline (100 µg/ml) than the vegetative cells (1 µg/ml) to reach similar GFP levels.

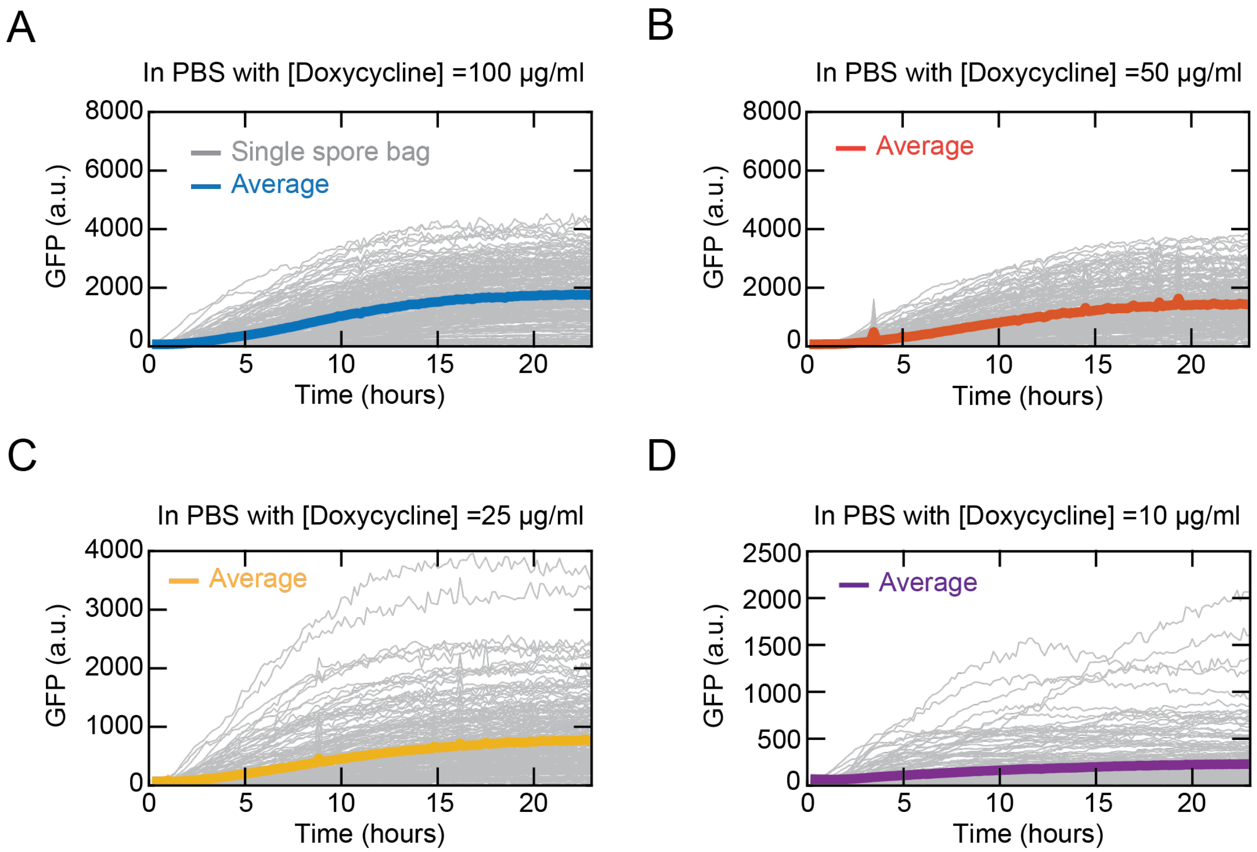


**Appendix Figure S12 - GFP expression without nutrients as function of time for different doxycycline concentrations (complements Fig. 3C).** GFP level of a single spore bag (grey curve) and population-level average (solid colored curves) over time while incubated in PBS (i.e., without any nutrients such as glucose and amino acids). We incubated the GFP-inducible spores in PBS with a set doxycycline-concentration (indicated above each graph) and then used a wide-field, epifluorescence microscope to measure the GFP levels of each spore bag for the next 22 hours. As seen in the grey curves plateauing over time, the GFP level of each spore bag reached a steady-state value by the end of the 22-hours of imaging. **(A)** [Doxycycline]=100 μg/ml, $n$ = 233 spore bags; **(B)** [Doxycycline] = 50 μg/ml, $n$ = 194 spore bags; **(C)** [Doxycycline] = 25 μg/ml, $n$ = 217 spore bags; **(D)** [Doxycycline] = 10 μg/ml, $n$ = 206 spore bags. Here, we measured the GFP levels continuously over time with a microscope for 22 hours to obtain the kinetics of GFP production whereas in Appendix Fig. S9, we performed one-time measurement on a microscope at the end of the 22 hours of incubation. We observed that spores could achieve higher GFP levels for the same doxycycline concentration if we incubated them in a continuously mixing liquid medium for 22 hours (Appendix Fig. S16) rather than in a stationary liquid medium inside a microscope well for 22 hours as shown here, due to the difference in culturing conditions.

**GFP-level remains nearly constant after removing doxycycine**

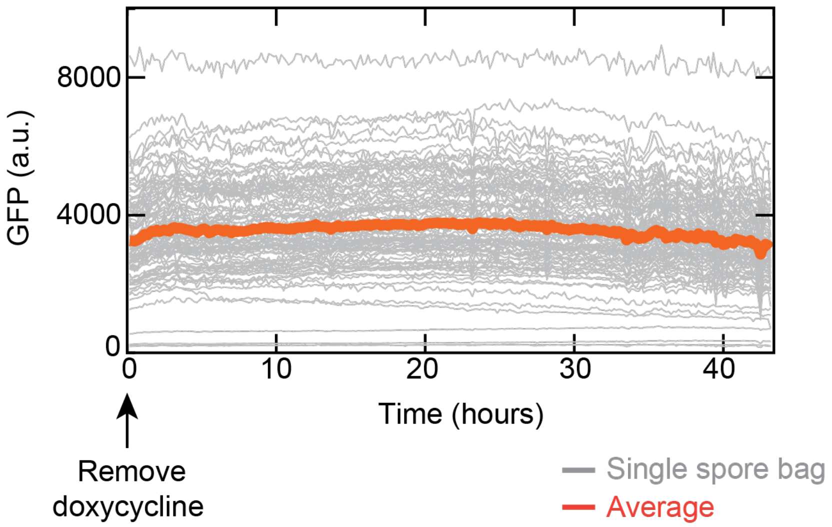


**Appendix Figure S13 - GFP level of a spore bag remains nearly constant after removing doxycycline (complements Fig. 3C).** GFP-levels of individual spore bags (grey curves) during 42 hours in PBS without doxycycline (*n* = 101 spore bags). The red curve shows the average GFP-level for these spore bags over time. We first induced GFP expression in these spores for 24 hours with 100 µg/ml of doxycycline. At the end of the 24-hour incubation, the GFP levels reached their steady-state values (Fig. 3C and Appendix Fig. S12). Then, we removed the doxycycline and washed away any residual doxycycline with PBS several times. We then incubated these spores in PBS at 30 $^{O}$C (start of this incubation marks "0 hours"). By doing so, we sought to understand why the GFP-levels reach steady-state values given that spores are not dividing to dilute away their accumulated copies of GFP - a vegetative cell's GFP level would reach a steady-state value because the production rate of GFP matching the dilution rate of GFP (dilution by cell divisions). If a spore bag's GFP level reached a steady-state value because of its GFP-production rate matching the GFP-degradation rate, then stopping the production of GFP by removing the doxycycline should cause decreases in its GFP-level. This is because GFP can then only degrade stochastically (thermally) and it cannot be replenished. As seen in the grey curves and the red curve, we did not observe any significant decreases in the GFP levels during the 42-hours that followed the removal of doxycycline. Thus, the reason that the GFP levels reached steady-state values is not because of the GFP-production rate matching the GFP-degradation rate during the 24 hours of induction with doxycycline. In fact, we see here that the GFP-degradation rate is nearly zero inside the spores. Thus, we can conclude that it is the eventual stopping of GFP-production, while doxycycline is still present, that causes the GFP-levels to reach steady-state values during the 24-hours of induction.

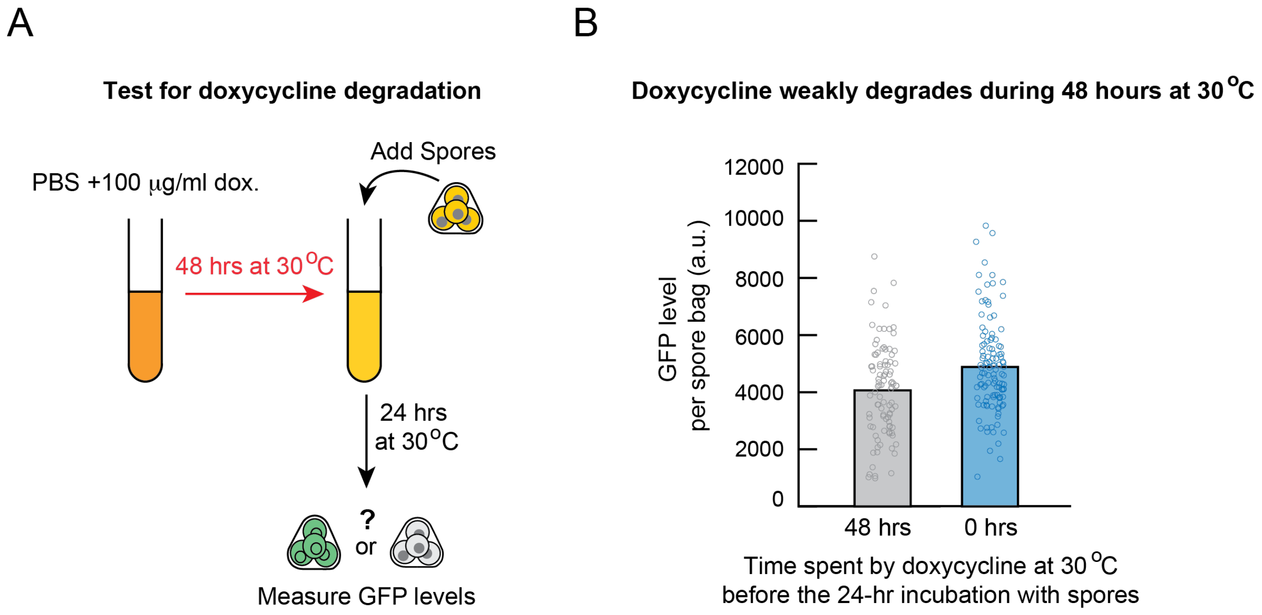


**Appendix Figure S14 - Doxycycline barely degrades during two days at 30 °C (complements Fig. 3C).** When we incubated spores in PBS with doxycycline, we always observed the GFP levels of individual spore bags plateauing after 15-to-20 hours (e.g., Appendix Fig S12). To test whether this is was due to large amounts of doxycycline degrading during after 15-to-20 hours, we first incubated PBS with 100 μg/ml of doxycycline for 48 hours in a rotating tube at 30 °C. Then, we incubated spores in this medium to induce GFP expression. After 24 hours of incubation, we measured the GFP levels in individual spore bags (grey data points, $n$ = 96 spore bags). As a control, we repeated the above steps but without first incubating the PBS containing the 100 μg/ml of doxycycline for two days (just as we did in all experiments involving GFP induction (Figs. 3-6)) (blue data points, $n$ = 116 spore bags). We found that, on average, incubating doxycycline for 48 hours caused the spores to express 20% less GFP (i.e., 4000 versus 5000 a.u.). Thus, doxycycline only weakly degrades after 48 hours. Hence, doxycycline degradation is not why the GFP levels plateau after 15-to-20 hours in Fig. 3C and any of the other GFP inductions in our work.

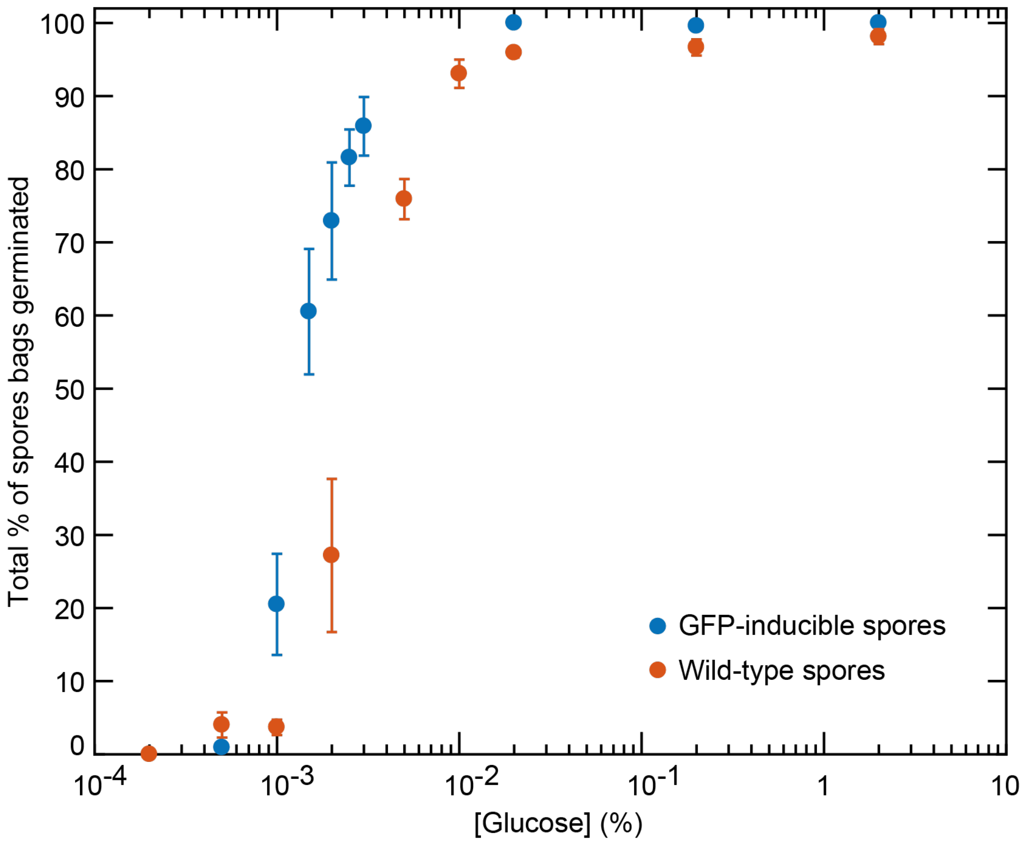


**Appendix Figure S15 - Comparing efficiency of germination for wild-type spores with that of GFP-inducible spores (complements Fig. 3D).** The GFP-inducible spores (Fig. 3A), aside from *GFP*, has additional genes (selection markers such as amino-acid biosynthesis genes) that we inserted during the yeast transformations (i.e., *ADE2, TRP1, URA3*). To see how these selection markers, as some of them pertain to amino-acid biosynthesis, might affect the percentage of spore bags that germinate for a given glucose-concentration, we gave different concentrations of glucose to the GFP-inducible spores (without any doxycycline, shown as blue data points) and compared the percentage of these spore bags that germinated with the percentage of wild-type spore bags that germinated (red data points) for the same glucose-concentration. As shown here, more GFP-inducible spores germinate than the wild-type spores for the same glucose concentration, but the overall trend is the same for both types of spores. $n$ = 3; error bars are s.e.m.

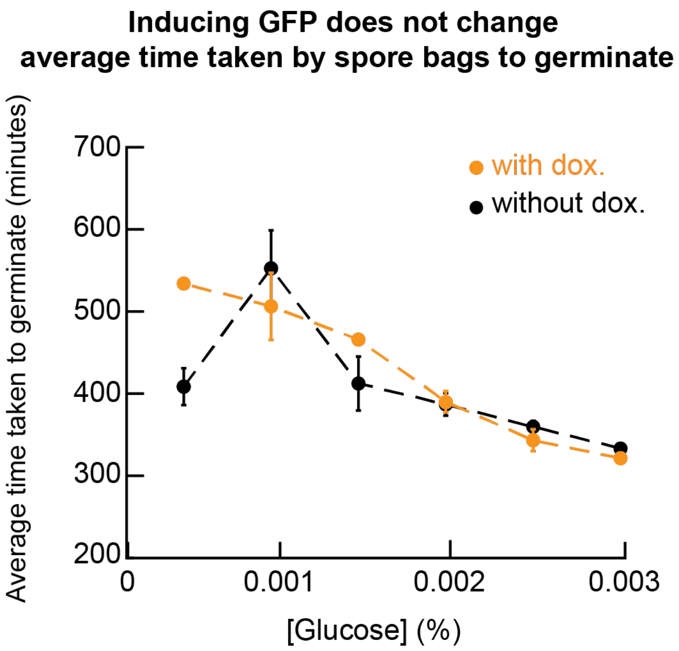


**Appendix Figure S16 - Inducing GFP production does not appreciably alter the average time taken by spore bags to germinate (complements Fig. 3D).** Same experiments and data as in Fig. 3D but now showing the average time taken by the GFP-inducible spore bags to germinate for low glucose-concentrations. Orange data points are for the GFP-inducible spores (Fig. 3A) that we first incubated with doxycycline for ~24 hours prior to receiving glucose (thus these spores have steady-state GFP levels prior to receiving glucose) and the black data points are for the GFP-inducible spores that did not receive any doxycycline (thus these spores have not produced GFP prior to receiving glucose). There is virtually no difference between the two conditions. Only for the lowest glucose concentration shown here, we see some differences in the average time taken for germination between the two conditions. This is due to small-number fluctuations (i.e., almost no spore bag germinates at such a low glucose concentration; we observed at most one or two spore bag germinating, if any, out of hundreds (c.f. Appendix Fig S1H)). Although not shown here, inducing GFP expression also does not appreciably alter the average time taken to germinate at much higher glucose concentrations than shown here (up to 2%-glucose) and it also does not appreciably alter the percentage of spore bags that germinate at each glucose concentration (shown in Fig. 3D). *n* = 3; error bars are s.e.m.

**Spore bags that can produce more GFP, without nutrients, are more likely to germinate for each glucose-concentration**

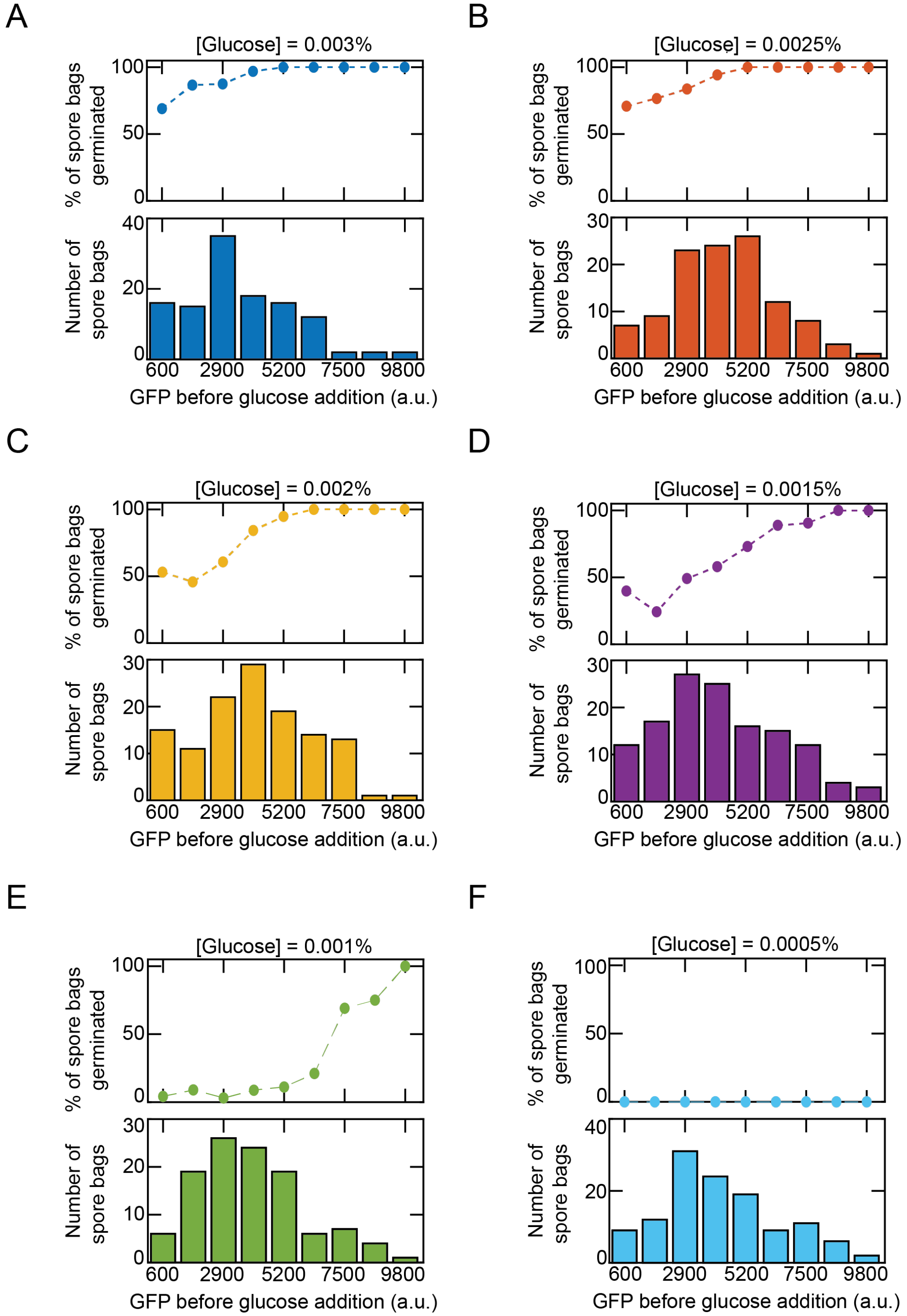


**Appendix Figure S17 - Spore bags that can express more GFP without any nutrients (higher GFP inducibility) are more likely to germinate for each glucose concentration**

**(complements Fig. 3E).** As liquid cultures inside rotating tubes, we incubated the GFP-inducible spores (Fig. 3A) in PBS first with 100 μg/ml of doxycycline for 22 hours. The spore bags' GFP levels reached steady-state values which we measured after transferring the spores onto microscope-imaging wells. After measuring the GFP levels at the end of the 22-hour incubation in this way, we removed the PBS containing doxycycline and then replaced it with a minimal medium that contained a relatively low concentration of glucose (indicated above each panel). We then measured, for each glucose concentration, how many of the spore bags that had similar GFP levels (i.e., GFP levels that fall within a binning range shown in the histograms above) germinated. **(A)** 0.003%-glucose, *n* = 118 spore bags; **(B)** 0.0025%-glucose, *n* = 113 spore bags. **(C)** 0.002%-glucose, *n* = 125 spore bags. **(D)** 0.0015%-glucose, *n* = 131 spore bags; (**E**) 0.001%-glucose, *n* = 112 spore bags; and **(F)** 0.0005%-glucose, *n* = 124 spore bags. The data shown here are for one of three biological replicates. We used all three biological replicates to construct the germination landscape in Fig. 3F. We used the procedure outlined here to measure all germination landscapes (Fig. 3F and in Fig. 4). We observed that spores could achieve higher GFP levels for the same doxycycline concentration if we incubated them in a continuously mixing liquid medium for 22 hours, as we did here, rather than in a stationary liquid medium inside a microscope well for 22 hours (Appendix Fig S12), due to the difference in culturing conditions. To be consistent, we used the method shown here (rotating liquid cultures) to obtain all germination landscapes and all the main conclusions in our study (e.g., probability of germinating as a function of GFP). We only cultured the GFP-inducible spores in microscope-imaging wells and continuously imaged them for 22 hours to show the kinetics of GFP-expression over time (only for Appendix Figs S10, S12, and S13 and Fig. 3C) but not for deriving the probabilities of germinating as a function of GFP levels. Importantly, our study's main conclusions, such as those expressing more GFP are more likely to germinate, are unaffected by the method of culturing GFP-inducible spores since these conclusions rely on relative levels of GFP rather than on the absolute levels of GFP.

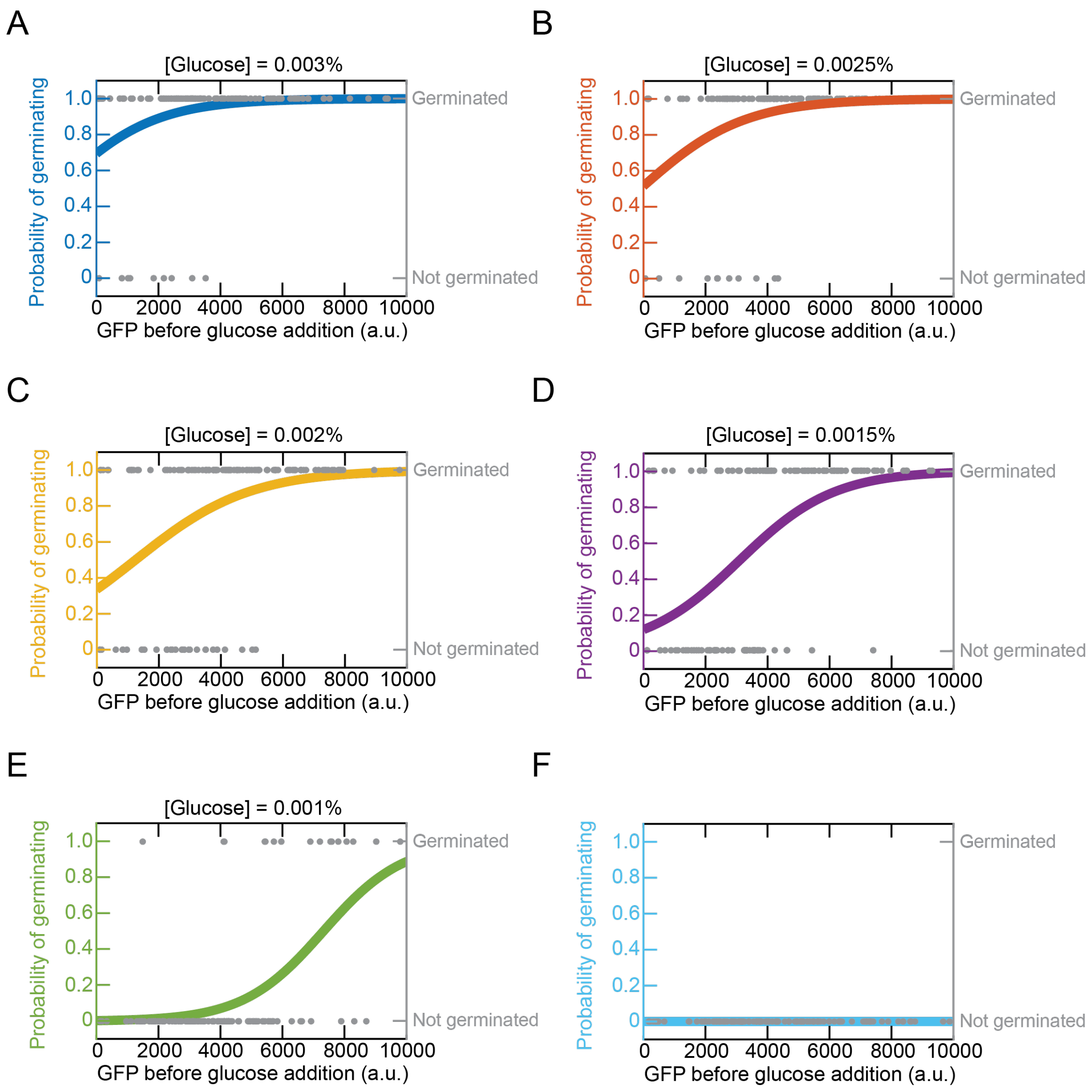


**Appendix Figure S18 - Statistical test with a logistic regression-fit establishes that spore bags that can express more GFP without nutrients (higher GFP inducibility) are more likely to germinate for each glucose-concentration (complements Appendix Fig S17 and the germination landscape in Fig. 3F).** Same experiments as in Appendix Fig S17 (i.e., GFP-inducible spores incubated for ~22 hours in PBS with 100 $\mu$g/ml of doxycycline, then measuring the steady-state GFP-levels of individual spore bags, and then finally adding glucose at concentrations as indicated above each panel (A-F) to determine which spore bags germinated). We used the same data as in Appendix Fig S17 except that now, we plot the data differently.

Each grey data point represents a single spore bag. For each spore bag, whose steady-state GFP-level we measured before adding glucose, we assigned it a "1" if it germinated or a "0" if it did not germinate after receiving the specified concentration of glucose. We then plot these grey data points (1 or 0) as a function of the GFP-level of each spore bag in the panels shown above (A-F). Afterwards, we performed a logistic regression on the grey data points, for each glucose-concentration, by fitting a logistic function, $p(x) = \frac{1}{1+e^{-(\beta_0+\beta_1 x)}}$, for the probability $p(x)$ that a spore bag with a steady-state GFP-level of $x$ germinates with a specified glucose-concentration. We used MATLAB's built-in “mnrfit” script to perform the logistic-regression fits (colored curves for each panel (A-F)). With the logistic function $p(x)$, testing a statistical link - that is, showing that there is a positive correlation between the GFP-level of a spore bag and its probability to germinate for a given glucose-concentration - is equivalent to testing whether the *x* (GFP-level before the spore bag receives glucose) is a sufficient predictor of the observed probability to germinate. We have done this by computing the p-value associated with the Wald test on the fit parameter $\beta_1$ which multiplies the *x* in $p(x)$. For every glucose-concentration, we found that the p-values were either below or equal to 0.01, meaning that the steady-state GFP-level of a spore bag indeed is a sufficient predictor for that spore bag's probability of germinating at the given glucose-concentration. Specifically, we found: **(A)** for a 0.003%-glucose: p-value $\approx$ 0.01, $\beta_1$ = -0.00063 $\pm$ 0.00049, *n* = 118 grey data points (92% germinated); **(B)** for a 0.0025%-glucose, p $\approx$ 0.002; $\beta_1$ = -0.0006 $\pm$ 0.00037, *n* = 113 grey data points (89% germinated); **(C)** for a 0.002%-glucose, p-value $\approx 3 \times 10^{-5}$; $\beta_1$ = -0.00054 $\pm$ 0.00026, *n* = 125 grey data points (77% germinated); **(D)** for a 0.0015%-glucose, p-value $\approx 4 \times 10^{-7}$; $\beta_1$ = -0.00066 $\pm$ 0.00025, *n* = 131 grey data points (61% germinated); **(E)** for a 0.001%-glucose, p-value $\approx 2 \times 10^{-5}$; $\beta_1$ = -0.00077 $\pm$ 0.00035, *n* = 118 grey data points (14% germinated); and **(F)** for a 0.005%-glucose, we did not observe any germinations in this data set. We have shown data from just one biological replicate here as a representative data set.

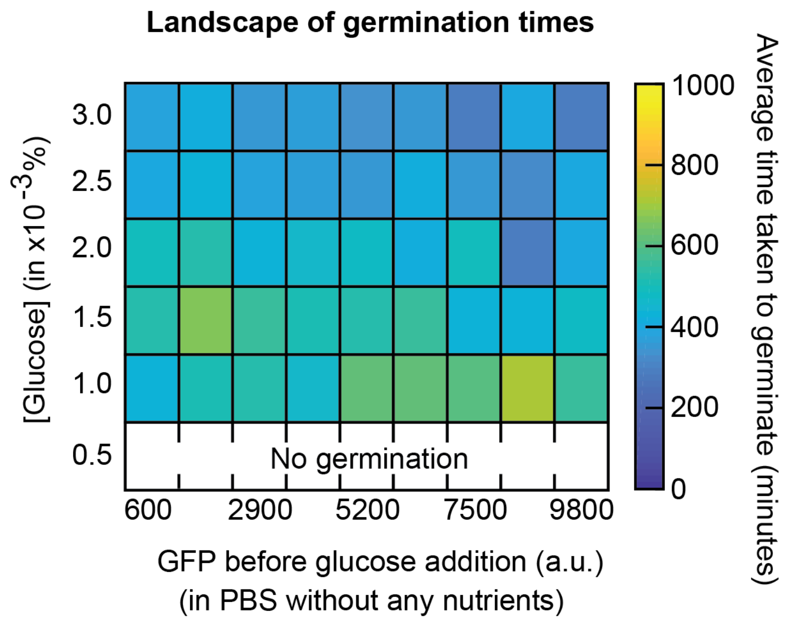


**Appendix Figure S19 - Average time taken to germinate depends weakly on GFP inducibility (complements Fig. 3F).** Each color represents the average time taken by a spore bag to germinate as a function of the glucose-concentration that it encounters and its steady-state GFP-level before it receives any glucose (result of GFP-induction with 100 μg/ml of doxycycline for 22 hours in PBS). Each color represents an average from three different populations of the GFP-inducible spores (from the same three biological replicate-populations as in Fig. 3F). For the lowest row, which represents a 0.0005%-glucose, the average time taken by a spore bag to germinate is undefined because we did not observe any spores germinating with this very low glucose-concentration. The lack of any dramatic changes in the shading of the colors across the pixels indicates that the average time taken by a spore bag to germinate depends weakly on the glucose concentration - a result that mirrors our earlier observation that the average time taken to germinate by the wild-type spores is also nearly independent of the glucose concentration (Fig. 1D). Crucially, we see here that the average time taken to germinate is nearly independent of a spore bag's steady-state GFP-level in PBS (as indicated by the absence of any clear changes in the shading of the colors across the pixels within a given row).

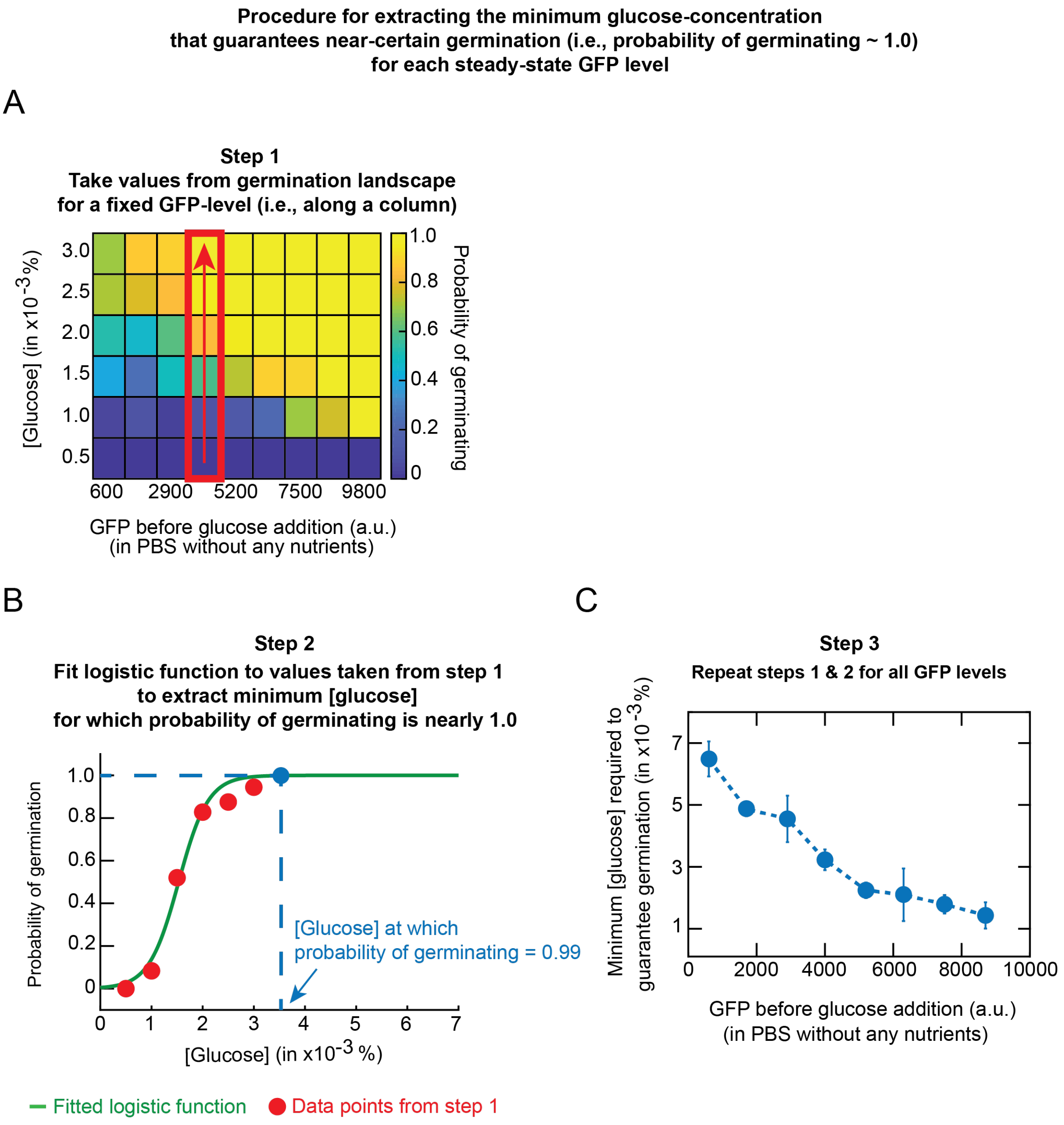


**Appendix Figure S20 - Three-step procedure shown here establishes that the minimum glucose concentration that is required for guaranteeing that a spore bag will germinate (i.e., probability of germinating ~ 0.99) decreases as the spore bag's GFP inducibility increases (complements Fig. 3F). (A)** The germination landscape (copy of Fig. 3F) groups the GFP-levels of spore bags into bins (columns of the heat map), with each bin (column) thus representing a defined range of GFP-levels. For each column of the germination landscape, we read-off the probability to germinate from each pixel (by moving up within the red box as shown in the figure). **(B)** We plot the values that we read-off from each pixel as a function of the glucose-concentration (red data points). We then fit a logistic function that has the same mathematical form as in Appendix Fig S18 but now with a different meaning: $p(x) = \frac{1}{1+e^{-(\beta_0+\beta_1 x)}}$. Here, $p(x)$ is the probability that a spore bag with the GFP-level specified in (A) germinates after encountering a glucose-concentration equal to $x$ (green curve) - note that the $x$ in Appendix Fig S18 represented a spore bag's steady-state GFP-level. Here we chose the logistic function for its simplicity. From the fitted logistic function (green curve), we can extract the value of $x$ (blue point) for which $p(x) \approx 0.99$ (i.e., the glucose-concentration for which the probability of germinating is 0.99). We chose 0.99 because choosing "1" will yield an artificially high value of $x$ given that the logistic function $p(x)$ asymptotically approaches 1 without ever reaching it. **(C)** Repeated the procedure in (A) and (B) for each column of the germination landscape yields the plot shown here: the minimum glucose-concentration that is required to guarantee that a spore bag will germinate (i.e., probability of germination ~ 0.99) if the spore bag's steady-state GFP-level in PBS is as specified in (A). The data points here are averages from three biological replicates (*n* = 3) and the error bars are s.e.m.

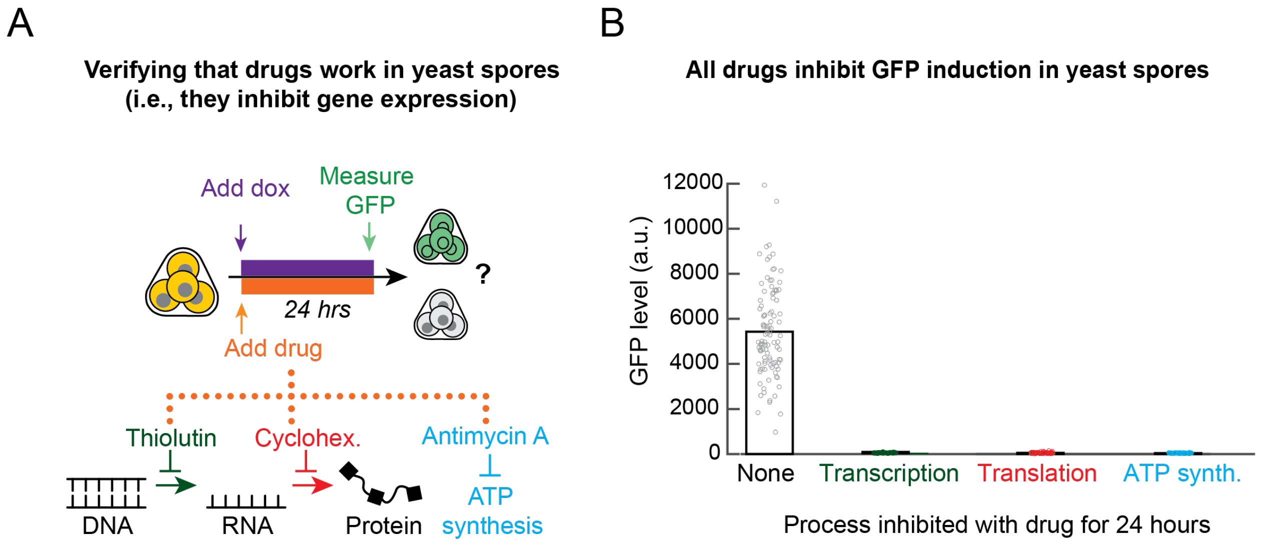


**Appendix Figure S21 - Thiolutin, cycloheximide, and antimycin A all function in yeast spores as shown by their complete inhibition of GFP induction (complements Figs. 4A-B).** **(A)** To verify that all three drugs that we used in Figs. 4A-B - thiolutin, cycloheximide, and antimycin A - actually functioned in yeast spores, we sought to test if all three drugs inhibited GFP expression in spores with the GFP-inducing synthetic circuit (Fig. 3A). We incubated spores for 24 hours in PBS with 100 μg/ml of doxycycline (to induce maximum possible GFP expression) and one of the three drugs - 10 μg/ml of thiolutin (inhibiting transcription) or 200 μg/ml of cycloheximide (inhibiting translation) or 100 μM of antimycin A (inhibiting ATP production). As a control, we also left out a drug in one case. **(B)** After the 24 hours of doxycycline with a drug, we measured the GFP levels of individual spore bags. Dots denote individual spores ($n$ = 62 for thiolutin, $n$ = 58 for cycloheximide, $n$ = 63 for antimycin A, $n$ = 75 for "None" (no drug)). The bars show the GFP-level averaged over all spores. The spores incubated in either thiolutin, cycloheximide, or antimycin A were not able to express any GFP. All three drugs thus inhibit gene expression in yeast spores.

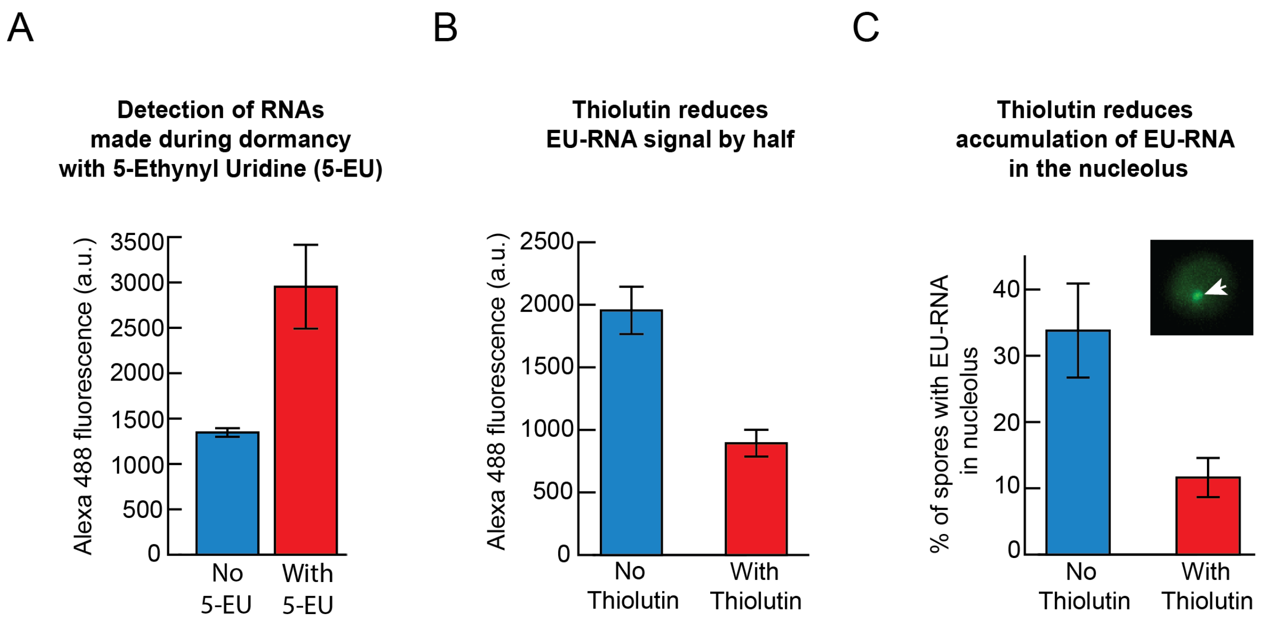


**Appendix Figure S22 - Detection of freshly made during dormancy with 5-Ethynyl Uridine (5-EU) (complements Figs. 4C-E). (A)** Alexa 488 fluorescence, on average, per spore bag after 24 hours of incubation in PBS without the 5-EU (blue) and with 1 mM of 5-EU (red). Hence, blue bar shows the background fluorescence which is ~2-fold lower than when 5-EU is present, indicating that the spores made RNAs during the 24-hour incubation. *n* = 3 populations (one of the populations is shown as histograms in Fig. 4D). **(B)** In order to further verify that the increase in fluorescence seen in (A) is from RNAs being synthesized during dormancy, we incubated spores for 24 hours in PBS with either 1 mM of 5-EU (blue) or 1 mM of 5-EU and 10 μg/ml of thiolutin (red). Thiolutin inhibits transcription. Thiolutin caused the Alexa 488 fluorescence per spore bag, on average, to be halved compared to the case without thiolutin. This is another validation that the 5-EU method for detecting freshly made RNAs is working in yeast spores. *n* = 3 populations. Errors bars are s.e.m. **(C)** (Blue) In ~35% of spores, we found RNAs being produced in the nucleoli of yeast spores that were incubated in PBS with 1 mM of 5-EU for 24 hours (blue). This number decreased to ~10% if the spores were in 10 μg/ml thiolutin with 5-EU for 24 hours (red). Inset picture shows a spore with the white arrow indicating the nucleolus where the RNA has accumulated during the 24 hours of incubation (images in Appendix Fig S23 actually prove that the spots like the one in the inset figure are RNAs in nucleoli). This indicates that thiolutin likely inhibits synthesis of non-coding RNAs as well. *n* = 3 replicates. Errors bars are s.e.m.

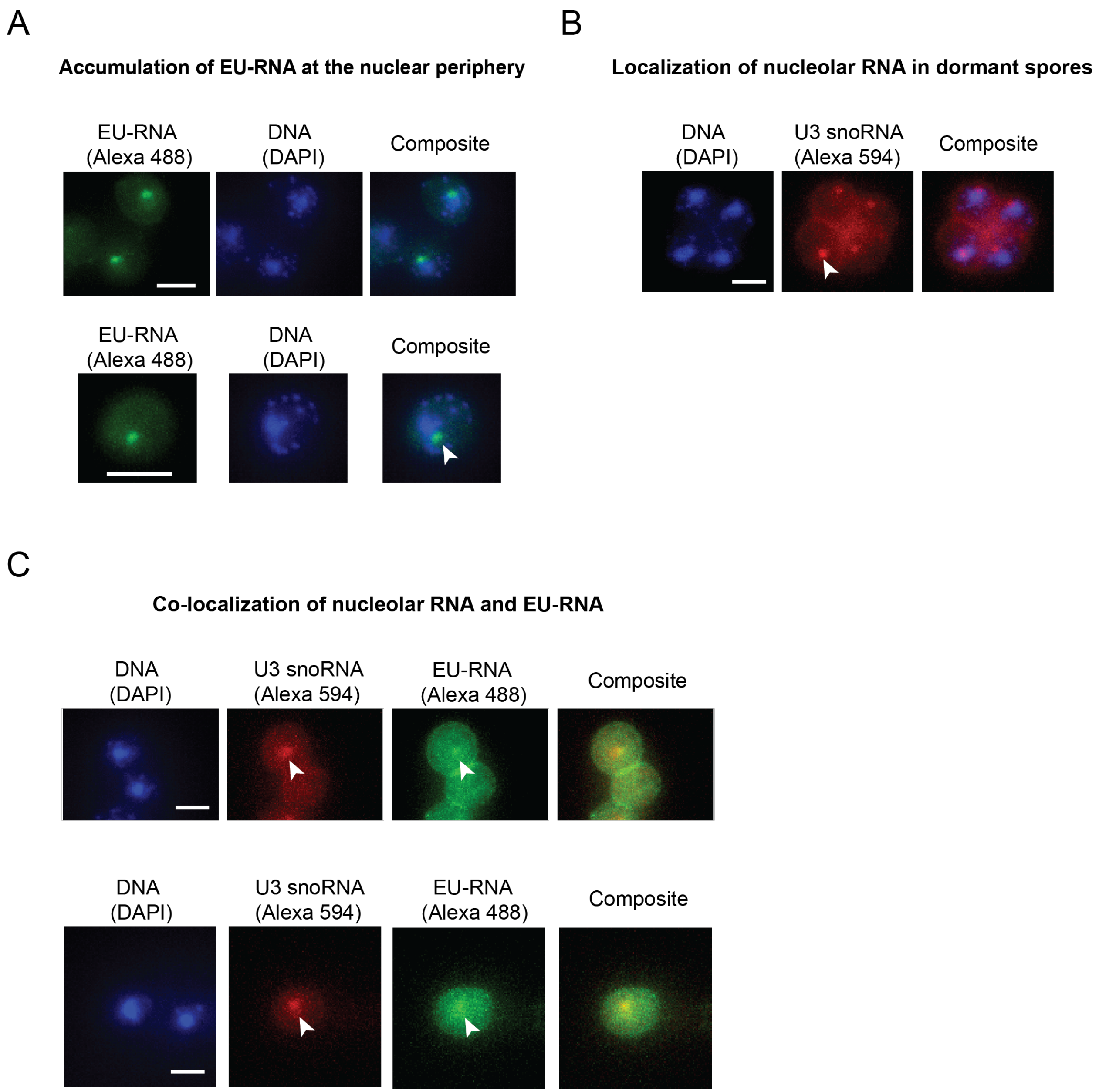


**Appendix Figure S23 - Synthesis of non-coding RNAs in spores' nucleoli (e.g., U3 snoRNA) detected with 5-Ethynyl Uridine (5-EU) (complements Figs. 4C-E). (A)** Images of fixed spores after 24 hours of incubation in PBS with 1 mM of 5-EU RNAs. During the fixation, we permeabilized the spores to let Alexa 488 fluorophores enter the spores and bind to the 5-EU-bound RNAs. Composite images show the merging of the Alexa 488 image (showing 5-EU bound

RNAs) and DAPI image (showing DNAs) of the same spores. Top row shows three spores and the bottom row shows one spore. For both rows, scale bar is 2 μm. White arrow indicates the single bright green fluorescence spot localized at the nuclear periphery. In vegetative yeasts, an image like the ones shown here represent RNAs accumulated in the nucleolus. **(B)** Shows four spores in a spore bag. Scale bar represents 2 μm. To confirm that the bright spots of Alexa 488 mark the nucleoli of spores, we used single-molecule RNA FISH to make U3 snoRNAs fluorescently visible (via Alexa 594 fluorophores). U3 snoRNAs localize in the nucleolus (Narayanan et al. *J. Cell Sci.* 2003). As in Narayanan et al., We used an anti-sense deoxyoligonucleotide probe against the yeast U3 snoRNA for single-molecule FISH whose sequence is ATTCAGTGGCTCTTTTGAAGAGTCAAAGAGTGACGATTCCTATAGAAATGA (purchased from IDT DNA). We used Alexa-594 fluorophore fused at the 3' end of the probe. We observed bright red fluorescence spots at the nuclear periphery (indicated by the white arrow), indicating the yeast spore's nucleolus. This result, with (A), indicate that the spores transcribe RNAs during dormancy. **(C)** To further confirm that dormant spores are making nucleolar RNAs during the 24-hours of incubation in 5-EU, we performed a sequential staining experiment in which spores were first incubated for 24 hours in PBS with 1mM of 5-EU. We then fixed the spores and performed single-molecule RNA FISH with the probe for the U3 snoRNA, using Alexa 594 as the fluorophore bound to the probe (as in (B)). Afterwards, we added the Alexa 488 fluorophores that would bind to the 5-EU labeled RNAs (as in (A)). Images here show the resulting spores. These images show clear red fluorescence spots (U3 snoRNAs in the nucleoli) and green spots. The green spots are relatively dim compared to the FISH spots likely due to the Alexa 488 fluorophores having less spaces to bind due to the Alexa 594 probes binding first to the same target RNAs. But even with the relatively dim 5-EU spots, we can observe that U3 snoRNA, indicating nucleolar RNA (red), co-localize with the 5-EU focal point in a spore. This result confirms that yeast spores produce the non-coding, nucleolar RNAs during dormancy. Scale bars represent 2 μm.

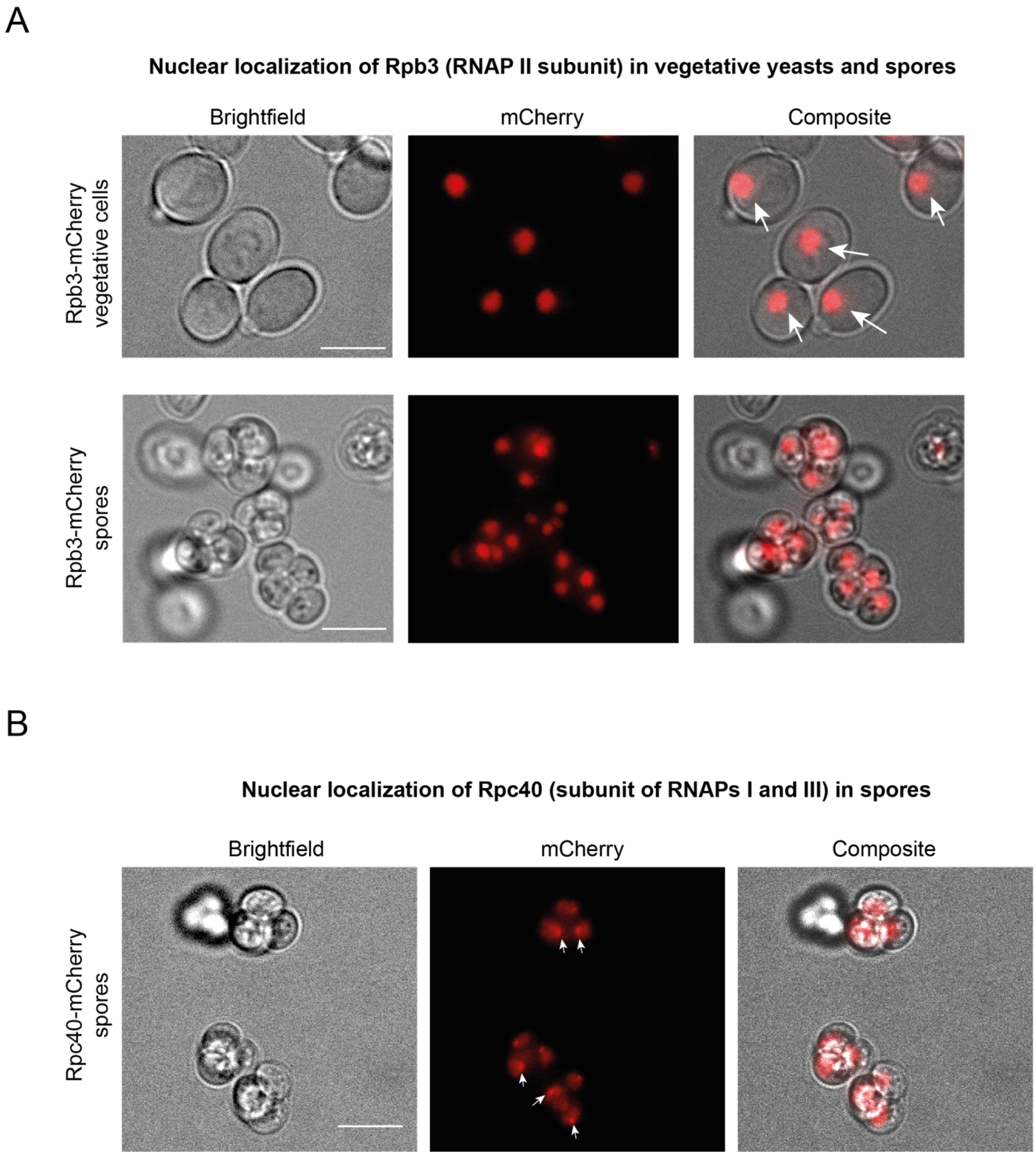


**Appendix Figure S24 - mCherry fluorescence is entirely confined to the nuclei of spores and vegetative yeasts with Rpb3-mCherry or Rpc40-mCherry (complements Figs. 4F-G). (A)** Representative images of diploid, vegetative yeasts (top row) and spores (bottom) whose Rpb3 (subunit of RNAP II) is fused to mCherry ("TS8" strain). As expected, mCherry fluorescence is almost entirely localized inside the nucleus of each spore or vegetative cell - indicated by one uniform red disk per spore or cell. White arrows show nuclear localized mCherry fluorescence. Scale bars represent 5 $\mu$m. **(B)** Representative images of spores with Rpc40 (subunit of both RNAPs I and III) fused to mCherry (“TS9” strain). White arrows show mCherry fluorescence localized at the nuclear periphery, as expected of RNAPs I and III since they are make mostly nucleolar transcripts.

**Spores take progressively longer times to germinate as time passes without nutrients**

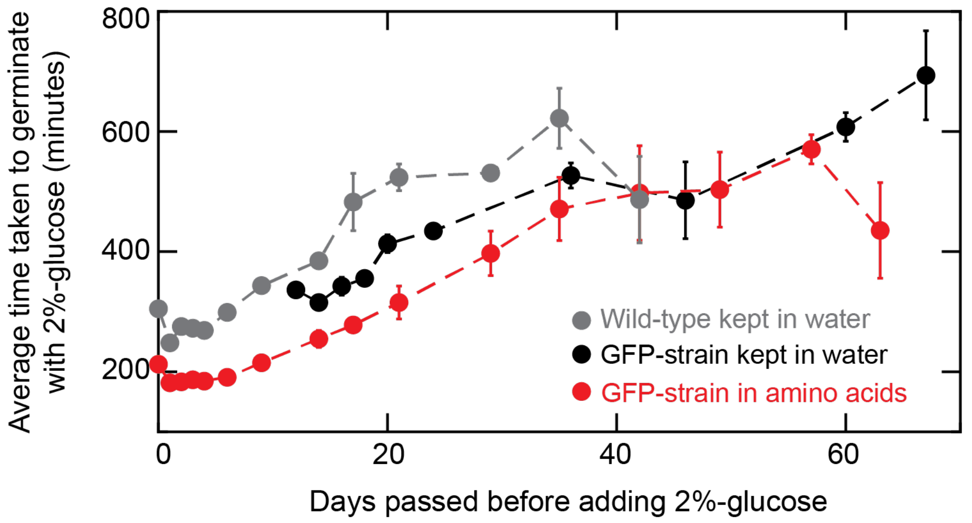


**Appendix Figure S25 - Dormant spores gradually take longer times to germinate as days pass by without any glucose (complements Fig. 5A).** Same experiments as in Fig. 5A but now plotting the average time taken by the GFP-inducible (black and red) and the wild-type (grey) spore bags to germinate due to a 2%-glucose, as a function of the number of days that they were incubated in either water (grey and black) or minimal media with essential amino acids (red) before receiving the 2%-glucose. In all cases, we that a spore bag, on average, requires more time to germinate the longer it is incubated in water or minimal media without any glucose. Importantly, we see that having the spores incubated in minimal media (which contains all the essential amino acids) does not majorly reduce the average time taken to germinate compared to having the spores incubated in water without any amino acids (compare red with black). Intriguingly, after 60 days (by which point almost all spores are dead according to Fig. 5A), the average time taken for germination is approximately 600 minutes, which is nearly equal to the average time taken by the wild-type spore bags to germinate on day 0 after encountering a 0.0005%-glucose - the smallest glucose-concentration for which we could observe germinations (see Fig. 1D). This suggests that both cases - one being a prolonged (~60-day) incubation without any nutrients and the other being a very low glucose-concentration - are probing spores at the "limits" of germination capabilities. *n* = 3; error bars are s.e.m.

**Protocol to distinguish dormant from dead spores by measuring their GFP levels without nutrients**

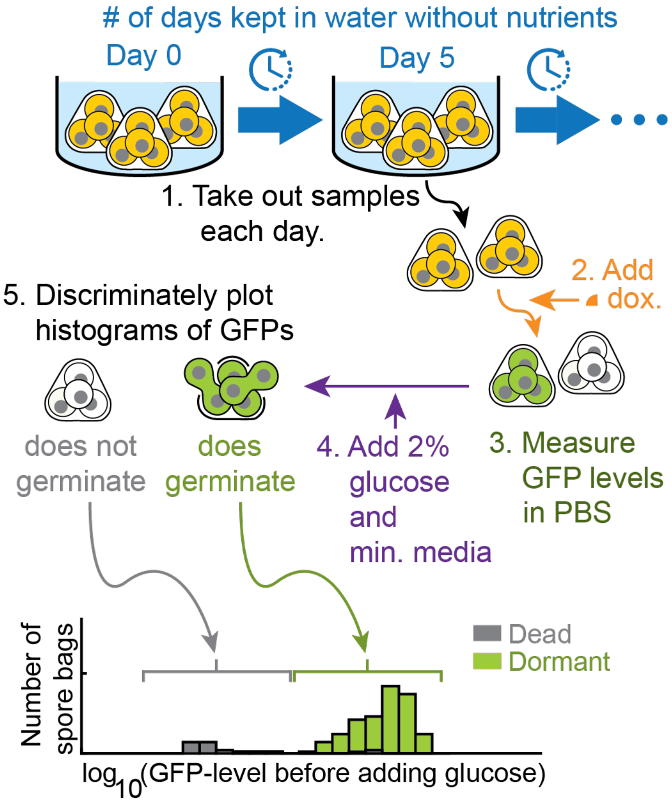


**Appendix Figure S26 - Protocol for Appendix Fig S27, which distinguishes dormant from dead spores by measuring their GFP levels without nutrients on different days of incubation in water without any nutrients at 30 $^{o}$C (complements Fig. 5B).** Protocol for measuring how the ability to express a generic gene (GFP) without nutrients changes over time in spore bags incubated in water for many days. The procedure shown here allows us to prove that spore bags lose their ability to express GFP (express a generic gene) and that as a result, the spore bag dies (i.e., once the ability-level reaches "zero"). There is a subtle procedural detail that we did not explain in the main text. By solely measuring the GFP-levels of individual spore bags after inducing them with doxycycline, we cannot distinguish between the values that correspond to dead or non-dead (still dormant) spores. This ambiguity is problematic because we want to demonstrate a cause-and-effect: the ability to express genes decreases *before* death. Since the GFP-level of dead spores is always close to zero (as defined by the relationship in Fig. 5B), by not excluding the GFP-level of dead spores in our analysis, we would then see that the mean GFP-level of the population would decrease over time, simply because spores are dying over time (Fig. 5A). Thus, after inducing GFP expression with doxycycline without nutrients, we add a 2%-glucose and then observe which spore bags germinate. We then plot the GFP-levels of the spore bags that germinate due to the 2%-glucose in the histograms in Appendix Fig S27 and discriminate them from the GFP-levels of dead spore-bags (the ones that do not germinate with the 2%-glucose). In other words, we plot the GFP levels of dormant spore bags in Appendix Fig S27 and discriminate them from the GFP levels of dead spore bags.

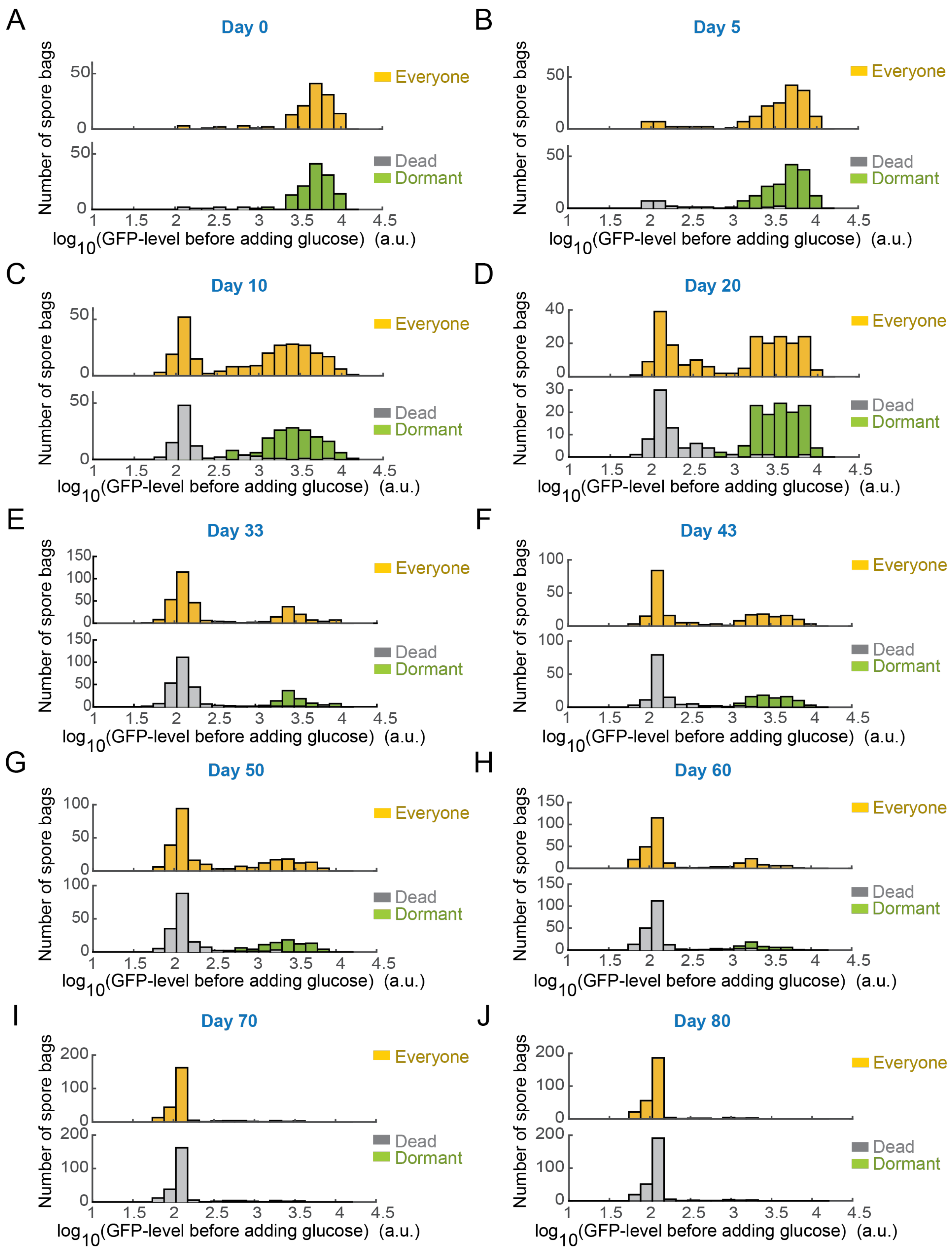


**Appendix Figure S27 | Dormant spores lose their gene-expressing ability before dying (complements Fig. 5B).** With the procedure in Appendix Fig S26, we measured the GFP-levels of each dormant spore-bag and dead spore-bag after incubating them in PBS with 100 µg/ml of doxycycline for 24 hours, on each of the incubation days in water without nutrients (from day 0 to day 80 of incubation in water without nutrients at 30 $^{O}$C). Yellow bars represent the entire population of spore bags; green bars represent dormant (alive) spore bags - these germinated after receiving a 2%-glucose; grey bars represent dead spores - these did not germinate after receiving a 2%-glucose. **(A)** Day 0: *n* = 225 spore bags in total (yellow bars), *n* = 221 dormant spore-bags (green bars); *n* = 14 dead spore-bags (grey bars). **(B)** Day 5: *n* = 180 spore bags in total (yellow bars), *n* = 158 dormant spore bags (green bars); *n* = 22 dead spore-bags (grey bars). **(C)** Day 10: *n* = 270 spore bags in total (yellow bars); *n* = 95 dormant spore-bags (green bars); *n* = 175 dead spore-bags (grey bars). **(D)** Day 20: *n* = 335 total spore-bags (yellow bars); *n* = 101 dormant spore-bags (green bars); *n* = 234 dead spore-bags (grey bars). **(E)** Day 33: *n* = 241 total spore-bags (yellow bars); *n* = 120 dormant spore-bags (green bars); *n* = 121 dead spore-bags (grey bars). **(F)** Day 43: *n* = 240 total spore-bags (yellow bars); *n* = 99 dormant spore-bags (green bars); *n* = 141 dead spore-bags (grey bars). **(G)** Day 50: *n* = 288 total spore-bags (yellow bars); *n* = 100 dormant spore-bags (green bars); *n* = 188 dead spore-bags (grey bars). **(H)** Day 60: *n* = 287 total spore-bags (yellow bars); *n* = 65 dormant spore-bags (green bars); *n* = 222 dead spore-bags (grey bars). **(I)** Day 70: *n* = 265 total spore-bags (yellow bars); *n* = 27 dormant spore-bags (green bars); *n* = 238 dead spore-bags (grey bars). **(J)** Day 80: *n* = 304 total spore-bags (yellow bars); *n* = 15 dormant spore-bags (green bars); *n* = 289 dead spore-bags (grey bars). By plotting the GFP-levels in base-10 logarithm, as we do here, we can see that the GFP levels of dead spore-bags (grey bars) are clearly distinct from those of dormant spore-bas (green bars).

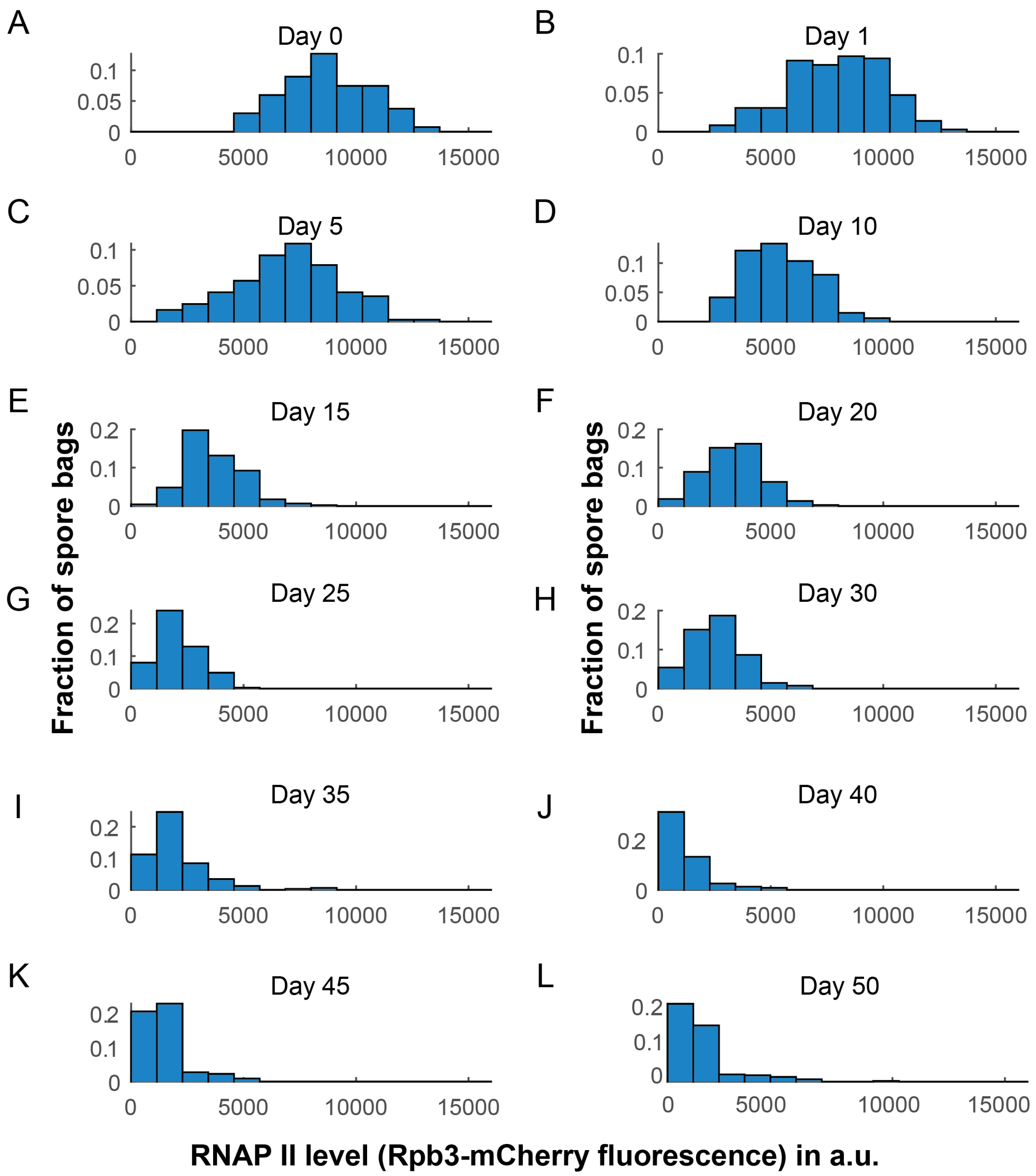


**Appendix Figure S28 - Histograms showing RNAP II levels of ageing spores on different days (complements Fig. 5C).** For each day of ageing shown in Fig. 5C, we show here a representative histogram (one population). Histograms show the RNAP II level of each spore bag in a population (i.e., fluorescence value from mCherry fused to Rpb3). Population averages of these histograms (3 histograms per day) are plotted in Fig. 5C. **(A)** Day 0. *n* = 134 spore bags. **(B)** Day 1. *n* = 181 spore bags. **(C)** Day 5. *n* = 184 spore bags. **(D)** Day 10. *n* = 169 spore bags. **(E)** Day 15. *n* = 228 spore bags. **(F)** Day 20. *n* = 191 spore bags. **(G)** Day 25. *n* = 194 spore bags. **(H)** Day 30. *n* = 139 spore bags. **(I)** Day 35. *n* = 160 spore bags. **(J)** Day 39. *n* = 117 spore bags. **(K)** Day 45. *n* = 109 spore bags. **(L)** Day 50. *n* = 196 spore bags.

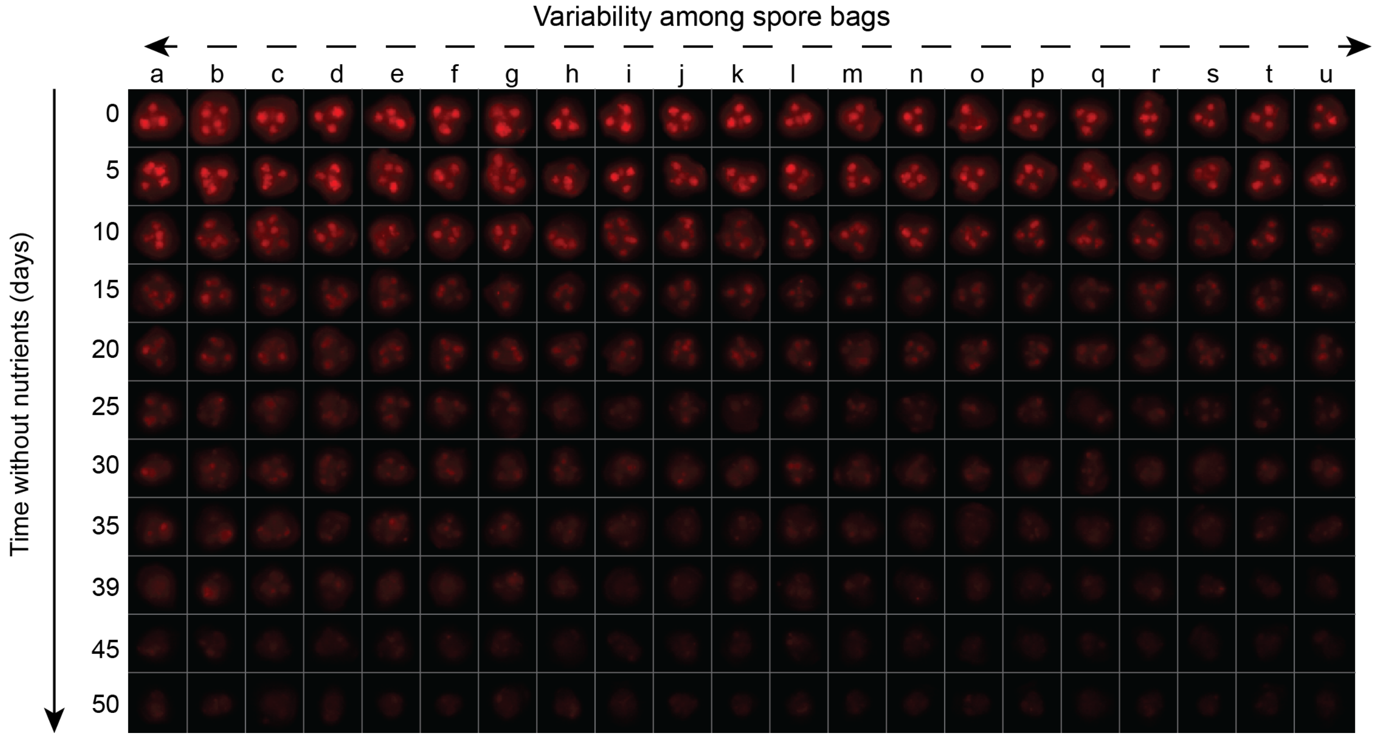


**Appendix Figure S29 - Montage of spore bags showing RNAP II level (Rbp3-mCherry fluorescence) on different days of ageing (rows), with multiple spore bags per row to show variability in RNAP II level among spore bags on the same day (complements Fig. 5C).** Each square contains a single spore bag and is 5 μm x 5 μm. Red color shows the mCherry fluorescence (Rpb3 fused to mCherry) and its intensity represents the amount of RNAP II in each spore bag. In each row, the spore bag becomes dimmer as one traverses towards the right end of each row (towards "u"). Each row displays 21 representative spore bags that were sampled on a given day, from a single population that was ageing in water for ~2 months at 30 $^{O}$C (the same rotating tube for ~2 months at 30 $^{O}$C) (data shown in Fig. 5C and Appendix Fig S28). Rows show different days of ageing (from 0 to 50 days). Since the RNAP II level is sorted in each row, from highest (a) to lowest (u), we can see the variability in RNAP II level among the spore bags on each day. This representation allows us to directly visualize the main phenomenon described in Fig. 5C: RNAP II level gradually decreases over days during dormancy (i.e., the pictures become dimmer within a column as we traverse from top to bottom).

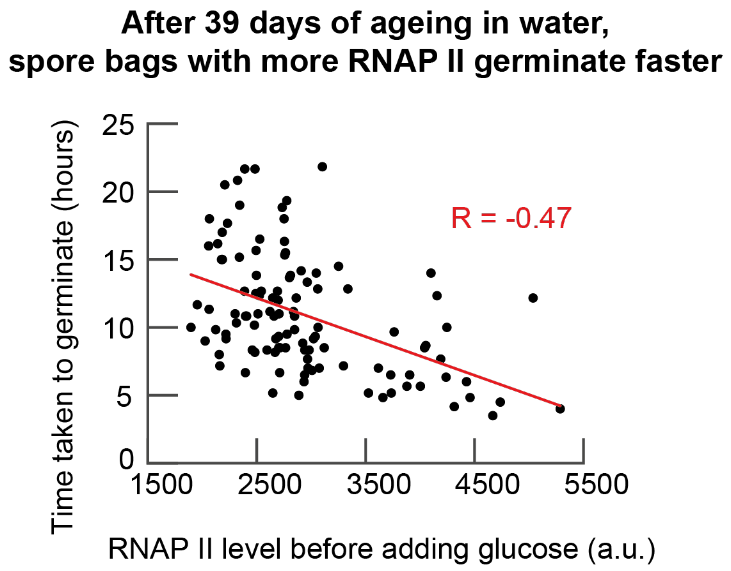


**Appendix Figure S30 - Having more RNAP II means germinating faster (complements Fig. 5F).** Time taken to germinate as a function of the RNAP II level (Rpb3-mCherry fluorescence level) in a spore bag. Each point represents a single spore bag from the same population. The population had been ageing in water for 39 days at 30 $^{O}$C (same data as Fig. 5F). Red line is a linear regression fit: R = -0.47, Pearson p-value = 1.67 x $10^{-7}$. *n* = 111 spore bags. Note that some germinate after more than 10 hours after receiving the 2%-glucose.

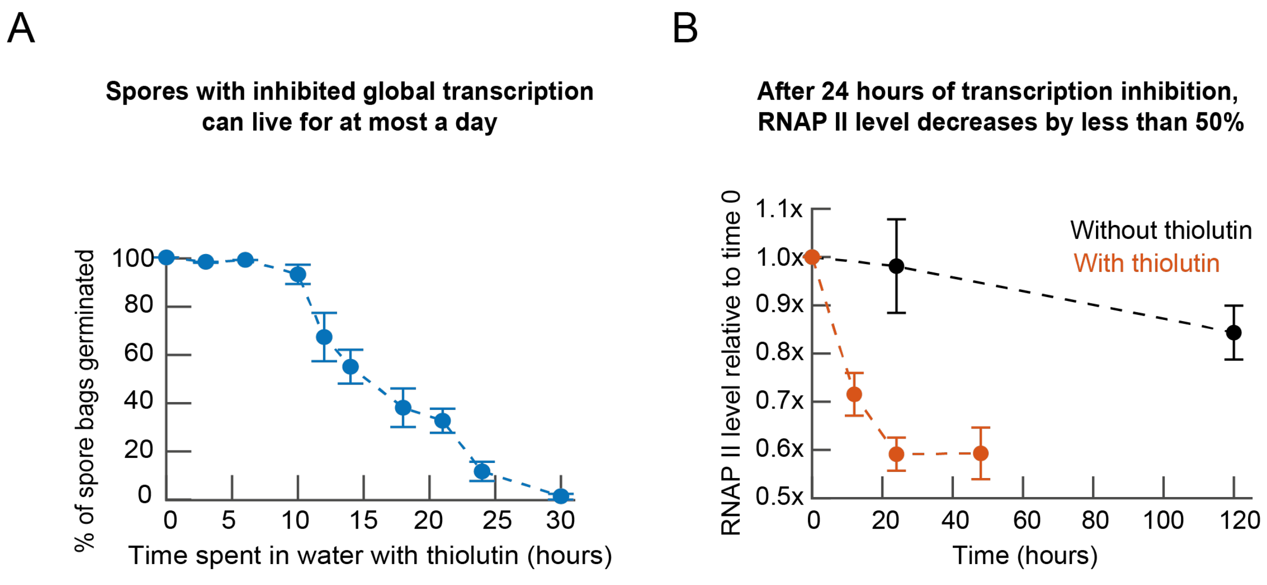


**Appendix Figure S31 - Spores with inhibited global transcription (with thiolutin) can live for at most a day while their RNAP II levels stay relatively high throughout (at most ~40% reduction in the RNAP II level after one day of thiolutin) (complements Figs. 7A-E). (A)** Experimental protocol shown in Fig. 7A. Percentage of spore bags germinated for different durations of incubation in water with 10 $\mu$g/ml of thiolutin. $n$ = 3 populations. Errors bars are s.e.m. **(B)** RNAP II level relative to its value at time zero (i.e., value at the start of incubation in water with thiolutin (red) or without thiolutin (black) for different amounts of time. $n$ = 3 populations. Errors bars are s.e.m. After 24 hours of thiolutin, almost all spores are dead (see (A)) while the RNAP II level in these dead spores is, on average, ~60% of the level at time zero. Moreover, RNAP II level did not continue to decrease (or decrease at the same fast rate) between 24 hours and 48 hours of thiolutin as seen by comparing the last two points.

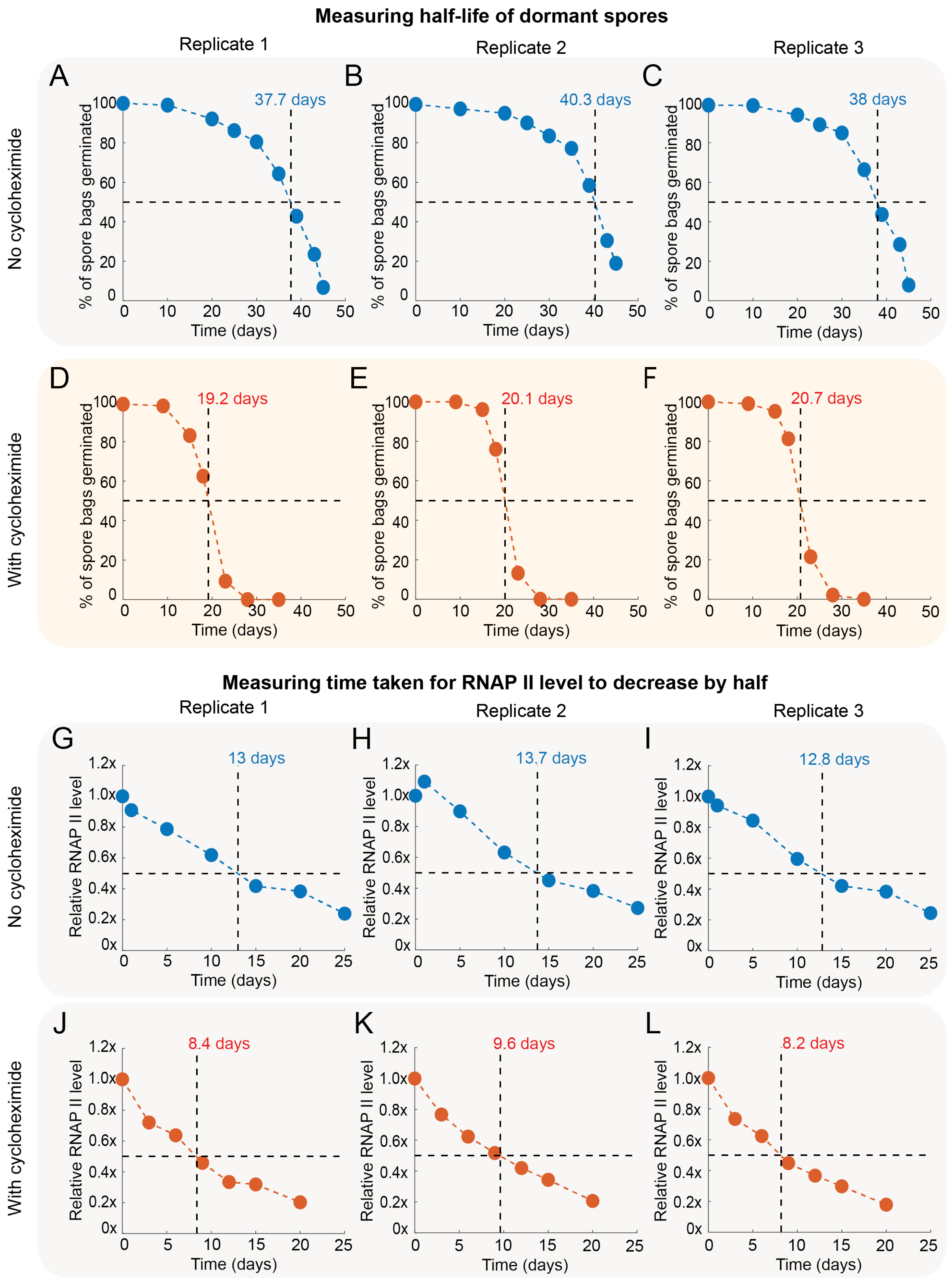


**Appendix Figure S32 - Estimating half-life of dormancy and RNAP II (complements Figs. 7A-E). (A-F)** Experimental protocol in Fig. 7A. Percentage of spore bags germinating after receiving a 2%-glucose after ageing in water without cycloheximide (blue points, A-C) or with 200 μg/ml of cycloheximide (red points, (D-F) for the designed days (values on horizontal axis). Each panel shows the half-life of dormancy (either in blue or red text) which we determined by a linear interpolation from finding the time at which 50% of the population had died (indicated by dashed black lines). **(G-L)** Average RNAP II level of dormant spore bags (Rpb3-mCherry fluorescence averaged over all dormant spore bags) relative to the level at the beginning of the ageing experiment in Fig. 7A. Blue points (G-I) are for spores aged in water without cycloheximide. Red points (J-L) are for spores aged in water with 200 μg/ml of cycloheximide. Each panel shows the time taken for the RNAP II level to decrease by 50% during ageing, which we determined by a linear interpolation, as indicated by the dashed black lines.

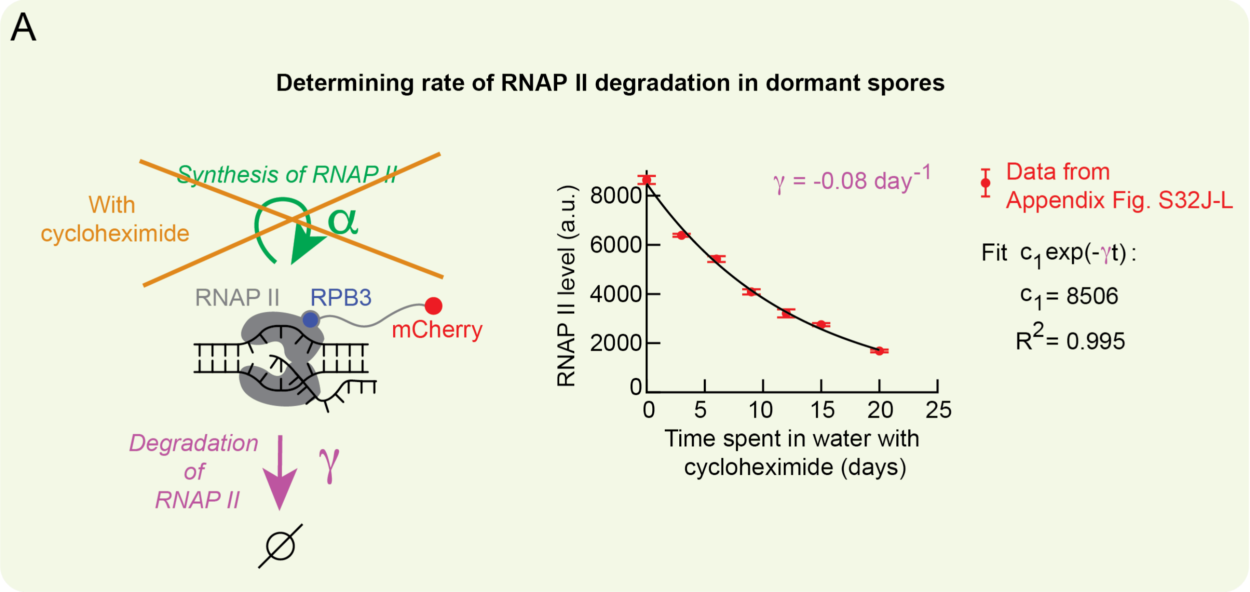


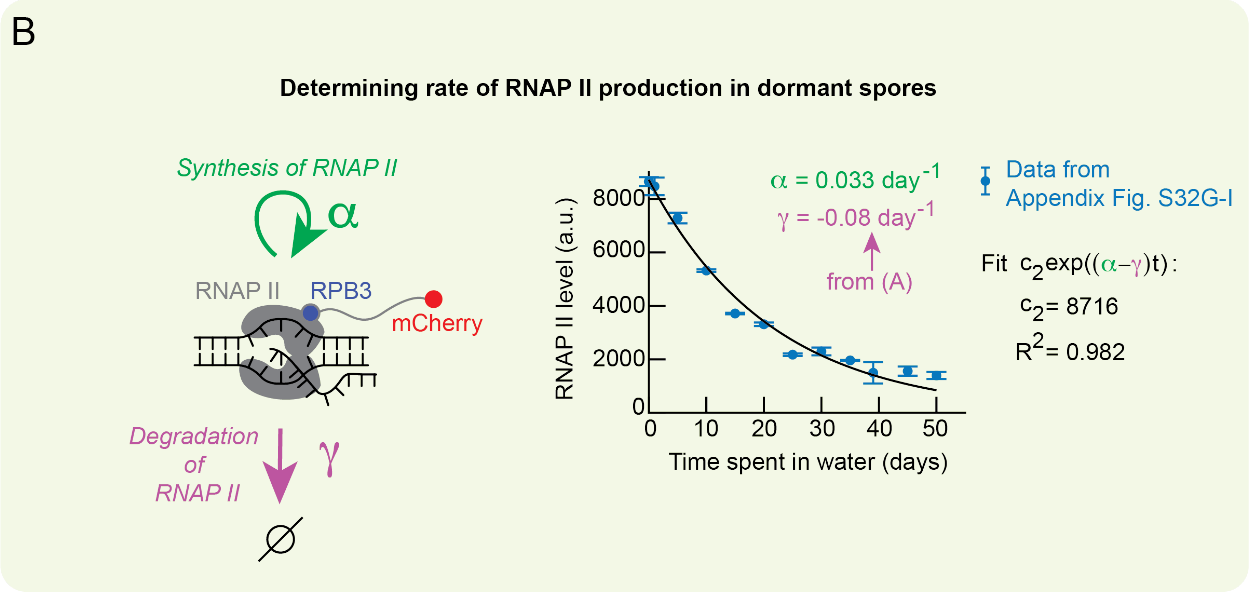


**Appendix Figure S33 - Estimating production rate of RNAP II during dormancy (complements Fig. 7F). (A)** We determined the degradation rate of RNAP II in dormant spores by using Rpb3-mCherry. For this, we assumed that when spores age in water with cycloheximide (translation inhibitor), their RNAP II level decreases purely due to the Rpb3 degrading, with each RNAP II molecule degrading with a probability per unit time of $\gamma$ (see cartoon). Hence, we have: $\frac{d[RNAP\ II]}{dt} = -\gamma.[RNAP\ II]$. Note that $\gamma$ is the degradation rate constant. This equation yields an exponentially decaying solution, $[RNAP\ II](t) = c_1.exp(-\gamma.t)$, where $c_1$ is initial RNAP II level. We fit the solution (black curve in the graph) to the data (red points in the graph) for the ageing experiment done with cycloheximide. Red data points are from averaging the three replicate populations shown in Appendix Fig. S32J-L; *n* = 3, errors bars are s.e.m. From the fit, we obtained a degradation rate constant, $\gamma = -0.08\ \mathrm{day}^{-1}$ with the fit agreeing well with the data ($R^2$=0.997). **(B)** Having determined the degradation rate, we now turn to determining the production rate of RNAP II during dormancy by using Rpb3-mCherry. For this, we assume that when spores age in water without any drugs, then the observed decrease in the RNAP II level during ageing occurs at a rate - a net loss rate - that is the difference between the production rate and degradation rate of RNAP II (with the degradation rate being higher than the production rate) (see cartoon). Then we have, $\frac{d[RNAP\ II]}{dt} = (\alpha - \gamma).[RNAP\ II]$, with $\gamma$ being the degradation rate determined in (A) and $\alpha$ being the production rate. We thus have as the solution, $[RNAP\ II](t) = c_2.exp((\alpha - \gamma).t)$ where $c_2$ is the initial RNAP II level at the start of the ageing. We fit the solution (black curve in the graph) to the data (blue points in the graph) for the ageing experiment without any drugs. Blue data points are from averaging the three replicate populations shown in Appendix Fig S32 G-I; *n* = 3, errors bars are s.e.m. Equivalently, we can fit the solution to the data in Fig. 5C. For the fit, we fix $\gamma = -0.08\ \mathrm{day}^{-1}$ as determined in (A). Hence $\alpha$ and $c_2$ are the only unknown parameters to fit. The fit yielded, $\alpha = -0.033\ \mathrm{day}^{-1}$, which agreed well with the data ($R^2$=0.982). If we ignore degradation by setting $\gamma = 0$, then we would get $[RNAP\ II](t) \propto exp(t/\tau)$, with $\tau = \frac{1}{\alpha} = 30.3\ \mathrm{days}$, being the "characteristic time" for RNAP II production in a dormant spore in water without any drugs. As a comparison, this is about 15-to-76-fold smaller than the characteristic time of 0.4-to-2 days measured in spores that have received a 2%-glucose and are on their way to germinating (Fig. 5E - blue points). These results establish that RNAP II production occurs during dormancy and that gene expression is occurring, albeit extremely slowly, during dormancy (e.g., expression of genes coding for RNAP II).

**Appendix Table S1 - List of genes for each transcriptional module (from Joseph-Strauss et al. *Genome Biol.* (2007)) (complements Fig. 2G and Appendix Fig S8).**

A transcriptional module is a group of genes that are involved in a related cellular process (e.g., stress response). An insightful work by Joseph-Strauss et al. (2007) has identified a list of transcriptional modules by studying yeast spores that germinated after receiving a saturating concentration (2%) of glucose. We performed transcriptome (RNA-seq) analysis of un-germinated spores that were primed by a low glucose-concentration (Fig. 2G and Appendix Fig S8). For this analysis, we used the transcriptional modules listed below - we looked at the specific genes listed below for each transcriptional module. This list is from Joseph-Strauss et al. *Genome Biol.* (2007).

| Transcriptional module | Genes in each module |
|---|---|
| Stress | YDL204W, TPS2, YGL037C, STF2, CTT1, SOL4, GRE3, OM45, YJR096W, YKL091C, TFS1, YLR251W, YLR252W, TSL1, YML128C, PGM2, YMR250W, YNL274C |
| Protein synthesis | RPL19A, RPS8A, RPL17A, RPS18B, RPS10A, URP1, CRY1, YS29B, RPL43A, YDL082W, YDL083C, SOS2, SOS1, RPS18A, RPS13C, RPL45, RPL15A, RP51B, YDR450W, RPL27B, RPL35B, RPL15B, RPS24EA, RPS8B, RPL17B, RPS26B, RPL32, RPL30A, RPL6A, CYH2, SUP44, SSM2, RPL9A, RPS31A, YGR034W, RPL16A, RPS28A, RPL30B, YST1, RPL14B, URP2, RPL4A, RPL27, MAK18, RPS7A, RPL5A, YIL052C, RPL13, UBI1, RPS25B, TIF2, YJL177W, RPS24A, RPS5, RPS7B, RPL14A, YKL056C, RPS27A, RPL17, RPS25, TIF1, UBI2, RPL4B, RPL13A, YST2, YLR061W, GRC5, UBI3, RPL35A, RPS33B, YLR325C, RPS31, YLR388W, RP10A, RPL16B, YML024W, YML026C, RP10B, YL16A, BEL1, YMR142C, YMR242C, RPL9B, RP23, YNL096C, RPL41A, RPS3, SSB2, RP28B, RPS16A, RPLA2, RPS21, RP28A, RPS16B, RPL25, TCM1, RPS30, RPS33A, RPL37B, YOR293W, RPL18A1, RPS12, EGD1, YPL079W, YPL090C, RPL37A, SSM1, YPR102C, RPS28B, YBR084CA, YER056CA, YFR031CA |
| Gluconeogenesis | YCR010C, ICL1, YFL030W, YGR067C, HXT5, YIL057C, MBR1, YKL187C, JEN1, PCK1, IDP2, FBP1, CYB2, YMR107W, YMR206W, MLS1, YNL194C, YNL195C, GAC1, LEE1, PXA1, YPR030W |

| | |
|---|---|
| Proteasome | PRE7, YBR062C, YBR173C, YTA5, RPN5, RPN4, YTA2, RPN8, PRE1, SUN2, PUP3, MPR1, YFR010W, PRE4, NIN1, SCL1, SUG1, UFD1, PRE9, PHB2, PUP2, ARC15, PRE3, CAP1, SBA1, YTA3, YKT6, YKR011C, YLR387C, YLR421C, GLO1, PRE8, PRE5, YNL155W, PRE6, CRL13, RPN7, PRE10, PRE2, RPN6 |
| Oxidative phosphorylation | PET9, COR1, ATP1, ATP3, YBR183W, YBR230C, ATP16, COX9, INH1, SDH4, ATP5, ATP17, QCR7, RIP1, YER053C, COX15, QCR6, COX4, COX13, CBP4, YGR182C, QCR9, COX6, QCR8, CYC1, MIR1, ATP2, ATP7, MDH1, HAP4, SDH3, SDH1, MCR1, SDH2, COX12, YLR294C, ATP14, COX8, NDI1, COX7, PBI2, COX5A, POR1, YNL100W, CIT1, CYT1, ATP4, ATP15, YPR020W, QCR2, YHR001WA |
| rRNA processing | YDR101C, YDR496C, ROK1, YGR103W, YGR145W, YGR245C, DRS1, PWP1, YLR222C, DBP9, YLR409C, YML093W, HAS1, YNL132W, YNL174W, YNL182C, YOR206W |
| Mating | PRM9, YAR033W, FUS3, YBL062W, FIG1, YBR156C, YBR158W, YBR223C, YBR225W, YBR226C, FUS1, KAR4, YCL074W, YCL075W, YCL076W, RVS161, FIG2, PCL2, RDI1, AFR1, YDR124W, ECM18, YDR241W, YDR249C, PAM1, YDR309C,YDR340W, STE14, MFA1, YER187W, STE2, YFL027C, YFL047W, AGA2, YGL052W, PRM8, IME4, YGL223C, GPA1, STE12, YHR097C, CHS7, PRM2, YIL060W, YIL080W, YIL082W, YIL083C, PRM5, YJL107C, PRM10, FAR1, ASG7, PGU1, GFA1, PGM1, PMU1, HYM1, YKL221W, KTR2, YLR042C, MID2, SST2, PRP39, PRM6, KAR5, CIK1, FUS2, YNL042W, MSG5, INP52, CHS1, YNL208W, PRM1, ERG24, AGA1, YOL095C, YOR129C, YOR343C, PRM4, PRM3, YPL193W, KAR3, YML048WA, YIL082WA, YMR304CA |
| Cell cycle: G1 | RFA1, SEN34, HTB2, HTA2, YBL009W, POL12, HHF1, HHT1, YBR070C, YBR071W, RDH54, RFC5, POL30, YBR089W, YCL022C, YCL024W, YCL061C, HCM1, MCD1, YDL018C, DUN1, YDL163W, CDC9, ASF2, MSH6, PDS1, HTB1, HTA1, YDR279W, GIN4, YDR528W, MNN1, PMI40, RNR1, RAD51, SSU81, ADK2, SMC1, CLB6, YGR151C, RSR1, YGR221C, YHR110W, YHR127W, SPO16, YHR154W, YHR173C, IRR1, YIL132C, SRO4, SMC3, HPR5, ASF1, RFA3, YJL181W, SWE1, POL32, PRI2, MIF2, HSL1, YKL108W, RAD27, YKR077W, YKR090W, KIM2, |

| | |
|---|---|
| | SPA2, YLL022C, STU2, YLR049C, CDC45, YLR183C, TUB4, SPH1, YOX1, OGG1, CTF18, YMR144W, SPT21, CLN1, HHF2, HHT2, PMS1, POL1, SPC98, YNL166C, BNI4, POL2, YIF1, TOF1, RFA2, YNR009W, YOL007C, YOL017W, MSH2, BUB3, DHS1, CDC21, YOR114W, YOR144C, NIP29, HHO1, RAD53, SVS1, IPL1, BBP1, CLN2, YPL267W, RLF2, CLB5, DPB2 |
| Cell cycle: G2-M | KIN3, YBL032W, CHS2, PHO3, CDC47, BUD3, YDR033W, SWI5, PMA1, ALK1, CDC20, DBF2, CLB1, WSC4, MOB1, YIL158W, YJL051W, BUD4, YKL130C, YLR084C, ACE2, YLR190W, YML034W, SUR7, YML058W, YML119W, CDC5, YMR032W, YMR215W, YNL057W, YNL058C, YOL070C, HST3, YOR315W, YPL141C, KIP2, IQG1, CLB2, YCR024CA |
